# The Molecular and Microbial Microenvironments in Chronically Diseased Lungs

**DOI:** 10.1101/676148

**Authors:** Alexey V. Melnik, Yoshiki Vázquez-Baeza, Alexander A. Aksenov, Embriette Hyde, Andrew C McAvoy, Mingxun Wang, Ricardo R. da Silva, Ivan Protsyuk, Jason V. Wu, Amina Bouslimani, Yan Wei Lim, Tal Luzzatto-Knaan, William Comstock, Robert A. Quinn, Richard Wong, Greg Humphrey, Gail Ackermann, Timothy Spivey, Sharon S. Brouha, Nuno Bandeira, Grace Y. Lin, Forest Rohwer, Douglas J. Conrad, Theodore Alexandrov, Rob Knight, Pieter C. Dorrestein, Neha Garg

**Affiliations:** Collaborative Mass Spectrometry Innovation Center, Skaggs School of Pharmacy and Pharmaceutical Sciences, University of California, San Diego, La Jolla, CA 92093, USA; Jacobs School of Engineering, University of California, San Diego, La Jolla, CA 92093, USA; Department of Pediatrics, University of California, San Diego, La Jolla, CA 92093, USA. Currently, Freelance Science Writer and Research Consultant; School of Chemistry and Biochemistry, Georgia Institute of Technology, Atlanta, GA 30332, USA; Department of Computer Science & Engineering, University of California, San Diego, La Jolla, CA 92093, USA; Structural and Computational Biology Unit, European Molecular Biology Laboratory, Heidelberg 69117, Germany; Biology Department, San Diego State University, San Diego, CA, USA; Department of Pathology, University of California, San Diego, La Jolla, CA 92093, USA; Department of Radiology, University of California, San Diego, La Jolla, CA 92093, USA; Department of Medicine, University of California at San Diego, La Jolla, CA 92093, USA; UC San Diego Center for Microbiome Innovation, University of California, San Diego, La Jolla, CA 92093, USA; Department of Bioengineering, University of California at San Diego, La Jolla, CA 92093, USA; Emory-Children’s Center for Cystic Fibrosis and Airways Disease Research, Atlanta, GA 30322, USA; Center for Microbial Dynamics and Infection, Georgia Institute of Technology, Atlanta, GA 30332, USA

## Abstract

To visualize the personalized distributions of pathogens, chemical environments including microbial metabolites, pharmaceuticals, and their metabolic products within and between human lungs afflicted with cystic fibrosis, we generated 3D microbiome and metabolome maps of six explanted lungs from three cystic fibrosis patients. These 3D spatial maps revealed that the chemical environments are variable between patients and within the lungs of each patient. Although the patients’ microbial ecosystems were defined by the dominant pathogen, their chemical diversity was not. Additionally, the chemical diversity between locales in lungs of the same individual sometimes exceeded inter-individual variation. Thus, the chemistry and microbiome of the explanted lungs appear to be not only personalized but also regiospecific. Previously undescribed analogs of microbial quinolones and antibiotic metabolites were also detected. Furthermore, mapping the chemical and microbial distributions allowed visualization of microbial community interactions, such as increased production of quorum sensing quinolones in locations where *Pseudomonas* was in contact with *Staphylococcus* and *Granulicatella*, consistent with *in vitro* observations of bacteria isolated from these patients. Visualization of microbe-metabolite associations within a host organ in early-stage CF disease in animal models will help elucidate a complex interplay between the presence of a given microbial structure, antibiotics, metabolism of antibiotics, microbial virulence factors, and host responses.

**Importance:** Microbial infections are now recognized to be polymicrobial and personalized in nature. A comprehensive analysis and understanding of the factors underlying the polymicrobial and personalized nature of infections remains limited, especially in the context of the host. By visualizing microbiomes and metabolomes of diseased human lungs, we describe how different the chemical environments are between hosts that are dominated by the same pathogen and how community interactions shape the chemical environment, or vice versa. We highlight that three-dimensional organ mapping are hypothesis building tools that allow us to design mechanistic studies aimed at addressing microbial responses to other microbes, the host, and pharmaceutical drugs.

## Introduction

An increasing rate of infection from multi-drug resistant opportunistic pathogens has become a significant burden in recent years. Proliferation of these pathogens due to overuse of antibiotics, including those of last resort (1–4), is a threat to human health and is already associated with increased mortality (5, 6). One reason for indiscriminate use of broad-spectrum antibiotics and combination therapy in complex polymicrobial infections is the lack of knowledge with regards to how microorganisms interact with each other, the host, and their chemical environment, leading to strategies that target bacterial pathogens broadly. Herein, specific microbial pathways that are involved in detrimental microbe-microbe interactions (7), microbe-host interactions (8) and microbe-drug interactions (9) can serve as new targets for targeted drug discovery. Thus, knowledge of such interaction-mediating microbial pathways and their prevalence will shape the future of drug discovery. In this regard, even though we have begun to appreciate the presence of multiple subpopulations by imaging community structures (10–12) and by genome sequencing (13–15), information about the specific microbial pathways in mediating the above mentioned interactions, molecular distribution of xenobiotic compounds, and how such distributions are associated with specific microbial structures within the context of a host is largely lacking.

We developed a methodology to map microbial and metabolite distributions in a human lung in three dimensions to identify pathways that may be mediating microbial interactions and to visualize the distribution of antibiotics in relation to microbial community structure (16). These three dimensional organ maps allow visualization of chemical and microbial microenvironments and consequently, may provide better insights into complex processes that take place within a host. Here we apply this methodology to elucidate spatial variation within and between the lungs of three individuals afflicted with cystic fibrosis (CF).

CF is genetic disease caused by a mutation in the cystic fibrosis transmembrane conductance regulator (CFTR) gene that results in defects of the encoded CFTR protein. The primary function of CFTR protein is an ion channel that regulates liquid volume (mucus) on the epithelial cells through secretion of chloride ions and inhibition of sodium absorption. Sticky mucus accumulates in the upper airways and lungs of CF patients, and serves as growth medium for various microbes, including opportunistic pathogens, resulting in chronic and recurrent polymicrobial infections. In the 1930s, the life expectancy for someone diagnosed with CF was only several years (17). Due to advances in modern medicine, including the use of antibiotics and better clinical management of the disease, individuals with CF can now expect to live on an average into their forties even though most patients are waitlisted for organ transplant by the time they reach adulthood (18). Improved clinical management is partly made possible by better understanding of the polymicrobial nature of the infections of the lung and development of antibiotic-based clearance of chronic infections targeting the polymicrobial community (19). However, the virulence of pathogens in microbial lung diseases such as CF, pneumonia, tuberculosis, and chronic obstructive pulmonary disease is mostly studied in cultures derived from pulmonary secretions and by genome sequencing, which does not represent complex *in vivo* conditions. The failure in treating an infection in a complex organ such as human lung may simply stem from the inability to treat a localized infection foci which can then spread to the entire organ or become systemic as, for example, in case of infections caused by *Burkholderia* (13, 14, 20, 21). Understanding how the production of microbial small molecules involved in pathogenicity and community interactions varies with lung biogeography leading to infection hot-spots will enable the development of targeted antimicrobials and improved drug delivery vehicles (14, 20, 22). Thus, CF presents an important test case for improving disease management strategies for polymicrobial infections, given better understanding of community structures and chemical environments within the host.

In this study, with patient consent, we mapped the chemical and microbial makeup of six explanted lungs, removed during surgery from three CF patients, by using 3D volume cartography to understand how microbes, microbial molecules, and medications are distributed and metabolized throughout the entire organ, providing insights into microbe-microbe interactions.

## Results and discussion

The explanted lungs of three patients afflicted with CF were sectioned to inventory and map the associated microbiome and metabolome in three dimensions onto lung models built from CT-scans acquired prior to surgery (Methods) (16). To perform 16S rRNA gene analysis, the tissue sections were swabbed, enabling detailed inventory of bacterial DNA present within the patients’ lungs (Table 1). We refer to our analysis of 16S rRNA gene as inventory of the bacterial DNA and not the bacteria themselves, since lungs associated with CF are known to contain a significant amount of DNA from dead cells as well as extracellular DNA (23). In total, six lungs from three patients contained bacteria that spanned 40 genera (Supplementary Table 1). Bar plots of the most frequently amplified genera and their relative abundances pooled for all anatomical locations are illustrated for each patient in Supplementary Figure 1. The relative abundances of these genera in individual sections of each patient is available through 3D maps (see below). The DNA of the most commonly occurring pathogenic organism in CF, *Pseudomonas aeruginosa*, was detected at highest frequency throughout the lungs of patients 1 and 3, whereas patient 2 was dominated by DNA from the emerging pathogen *Stenotrophomonas*. Even though the microbial population within CF-associated lungs can be heterogeneous (24), dominance of a single pathogen at end-stage CF disease has been described extensively in previous studies (25–27).

**Table 1.**
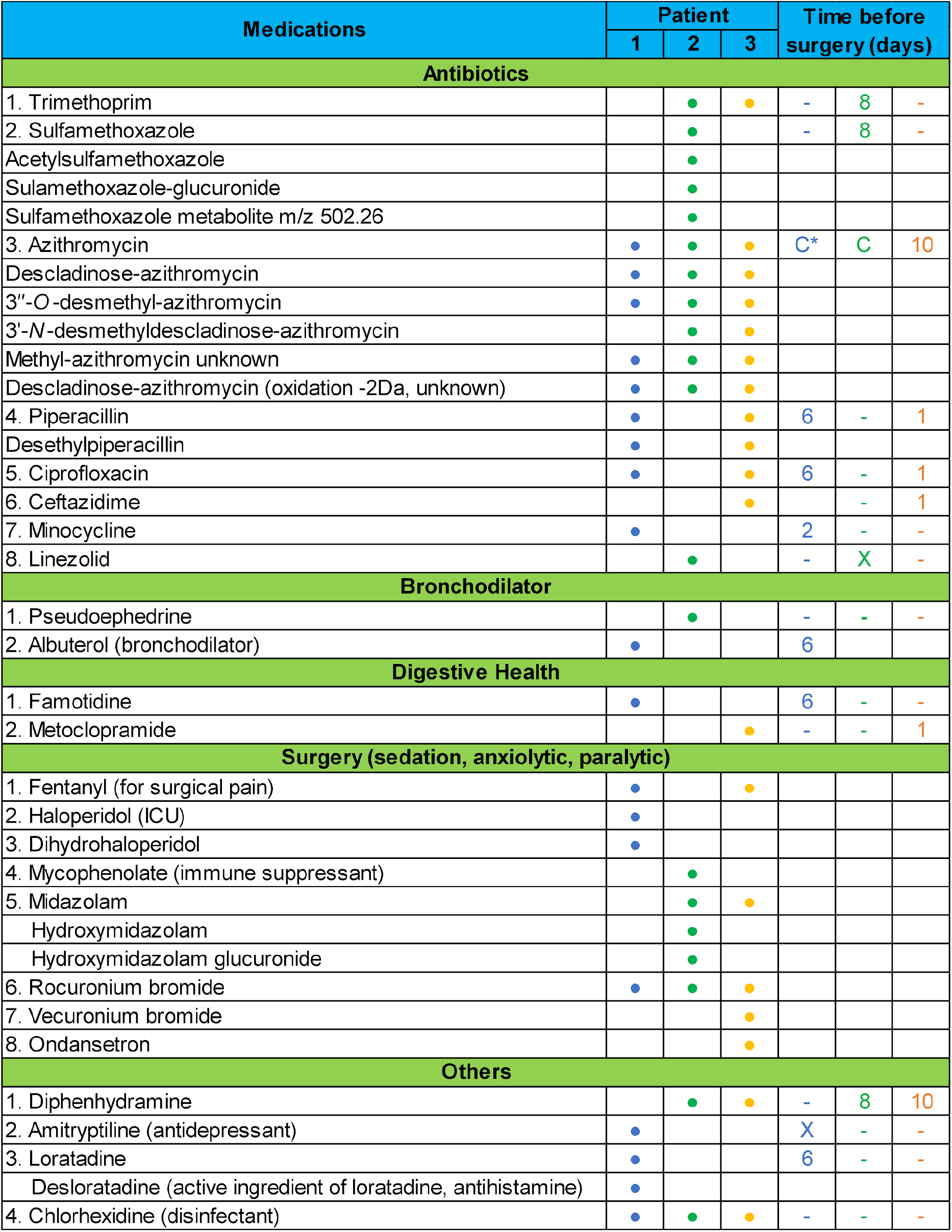
The medications detected in the MS data and time of administration prior to the day of lung explantation surgery. C refers to: “continuously administered”

The PCoA analysis of microbiome data with weighted UniFrac distance showed clustering between both lungs of patient 1 and the right lung of patient 3 along the first two principal axes (Supplementary Figure 2a). Samples from the left lung of patient 3 clustered separately and the comparison of 10-most abundant OTUs further highlighted the differences between the microbial communities present in the left lung of patient 3 (Supplementary Figure 2b and Supplementary Figure 3). Apart from the dominant pathogen, the overall microbiome between and within patients was different along the second and third axis (Supplementary Figure 2 c,d). The Canberra distance metric yielded less apparent but visible separation of patients’ microbiome data in PCoA space (Figure 1 a,b); in some cases this variation within patient’s own lungs was found to be greater than between patients (Figure 1b). A more homogeneous clustering patterns is observed in the 16S data when the unweighted UniFrac metric, a phylogenetically informed metric, was applied (Supplementary Figure 4). The differences arise due to the qualitative nature of unweighted UniFrac and the quantitative comparisons possible with the Canberra metric.

**Figure 1.**
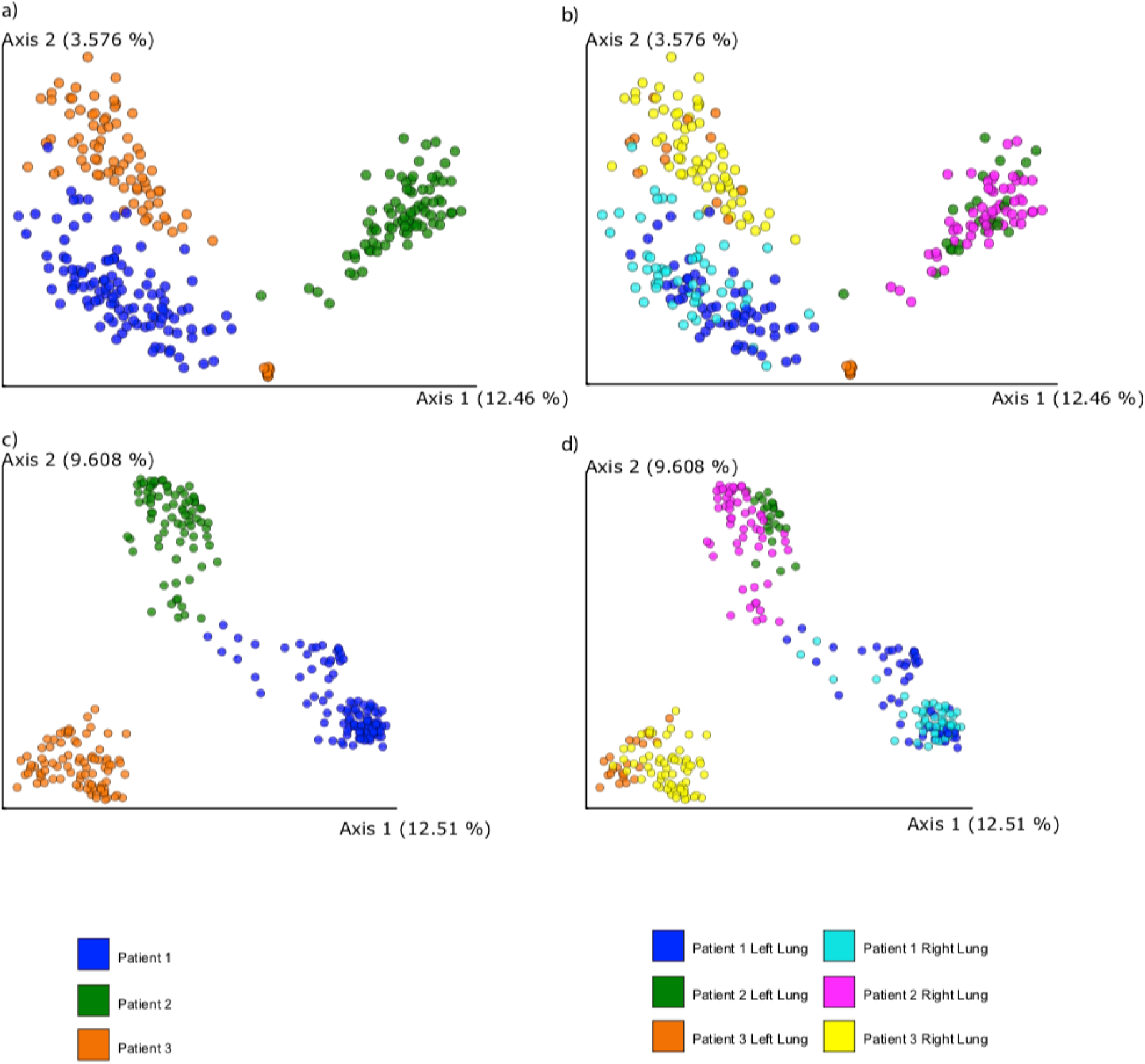
Principal coordinate plot of metabolome and microbiome from lungs of three patients in the study. Visualization was performed in the Emperor software using Canberra distance metric (28). a), b) PCoA plots of 16S rRNA sequencing and c), d) mass spectrometry data, respectively.

To map the relative frequency of microbes onto the 3D lung models, we used our previously described methodology (16). The distribution of prevalent (*Pseudomonas* and *Staphylococcus*) and emerging (*Stenotrophomonas* and *Achromobacter*) microbes in the CF-associated lungs is displayed in Figure 2. Although nearly uniform distribution of dominant pathogens (*Pseudomonas* in patients 1 and 3, *Stenotrophomonas* in patient 2) was observed, all other microbes were distributed unevenly, often relegated to niche spots. For example, *Achromobacter* was mainly localized in the apex of the right lung of patient 3 while *Staphylococcus* was present in the lower lobe of both lungs of patient 1, at the apex of the lungs of patient 2 and in the middle and lower lobes of the lungs of patient 3. The dominant pathogen, *Stenotrophomonas*, showed uniform distribution in the lungs of the patient 2 and differential distributions in the lungs of patients 1 and 3 (Figure 2). A degree of stratification is expected based on the availability of oxygen: *Achromobacter* and *Stenotrophomonas* are strict aerobes whereas *Staphylococcus* and *Pseudomonas* are facultative anaerobes residing as biofilms in airway mucus of CF patients with the potential of undergoing anaerobic metabolism (29). Furthermore, patients 1 and 3 not only share the dominant pathogen, their microbial communities are also more similar to each other than either is to that of the patient 2 (Supplementary Figure 3 and 5). Selection pressures from competing microbes and chemical microenvironments including antibiotic distributions, further leads to stratification of niches occupied by specific organisms. To compare the microbial and chemical environments, we next annotated the mass spectrometry data acquired on the tissue sections and mapped it onto the 3D models of the lungs of these patients (see below).

**Figure 2.**
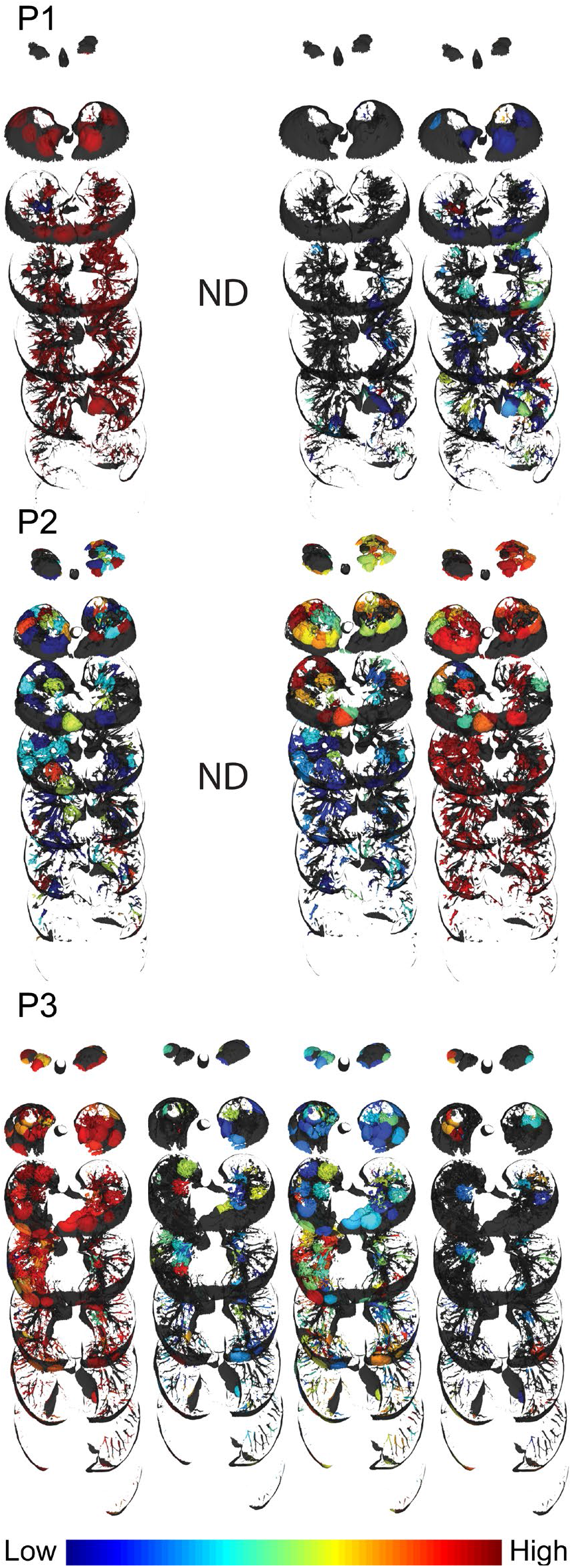
Distribution of microorganisms. Distribution of *Pseudomonas, Achromobacter*, *Staphylococcus*, and *Stenotrophomonas* (right to left) are shown for all three patients. P1 - patient one, P2 - patient two and P3 - patient three, ND - not detected. Intensity scale is provided at the bottom right. Full visualizations of microbial maps can be accessed via links: <u>patient 1</u>, <u>patient 2</u> and <u>patient 3</u>.

To annotate molecular ions detected using high resolution mass spectrometry (MS)-based untargeted approach, molecular network analysis was performed using the Global Natural Product Social Molecular Networking (GNPS) infrastructure (30). Molecular networking allows for reduction and organization of the overwhelming amount of chemical information generated (in terms of mass spectra) in a high-resolution untargeted MS approach. The data reduction is performed by combining and displaying the identical MS/MS spectra as a single node and by displaying similar spectra as connected nodes (30, 31). Similarities in MS/MS spectra relate to similarities in chemical structures, so oftentimes such connected nodes represent chemical and biological transformations of a molecule. In this study, 676,451 MS/MS spectra were filtered and merged into consensus spectra, producing 9,874 nodes (Figure 3a). The patient-specific molecules were displayed by assigning a specific color to each patient in the molecular network analysis (Figure 3a). In addition to annotating known compounds, molecular networking revealed related molecules that differ by oxidation, methylation, acetylation, hydroxylation, glycosylation, chain length, and saturation of alkyl chains, which enabled identification of previously undescribed metabolites of administered pharmaceuticals and microbial quinolones, as described below for azithromycin and *Pseudomonas aeruginosa* quinolones. The frequency of detection of the antibiotics across patients’ samples is shown in Figure 3b, with corresponding clusters from the full network displayed for each antibiotic. The nodes in antibiotic cluster represent the metabolic transformations of the antibiotic. Thus, molecular networking provides a glimpse into metabolic processes. The resulting molecular network revealed that among three patients, remarkably, only about 27.6% of detected molecular features were shared, highlighting the diversity of chemistry present in diseased human lungs (Figure 3c). All three patients in this study had different mutations in the CFTR gene (see Materials and Methods) and patients 2 and 3 were also diagnosed with CF-related diabetes. Two of the three patients suffered from chronic infections by *Pseudomonas aeruginosa*. Thus, various factors may play a role leading to the observed chemical diversity, which may arise from microbial (e.g., virulence and quorum sensing metabolites such as quinolones), host (e.g., bile acids, amino acids, sugars, eukaryotic lipids, fatty acids, sterols, peptides, immune-related molecules), and xenobiotic molecules. The diversity of these metabolites in CF sputum has been previously characterized and many of the same compounds were also found in the lung tissue in this study (32).

**Figure 3.**
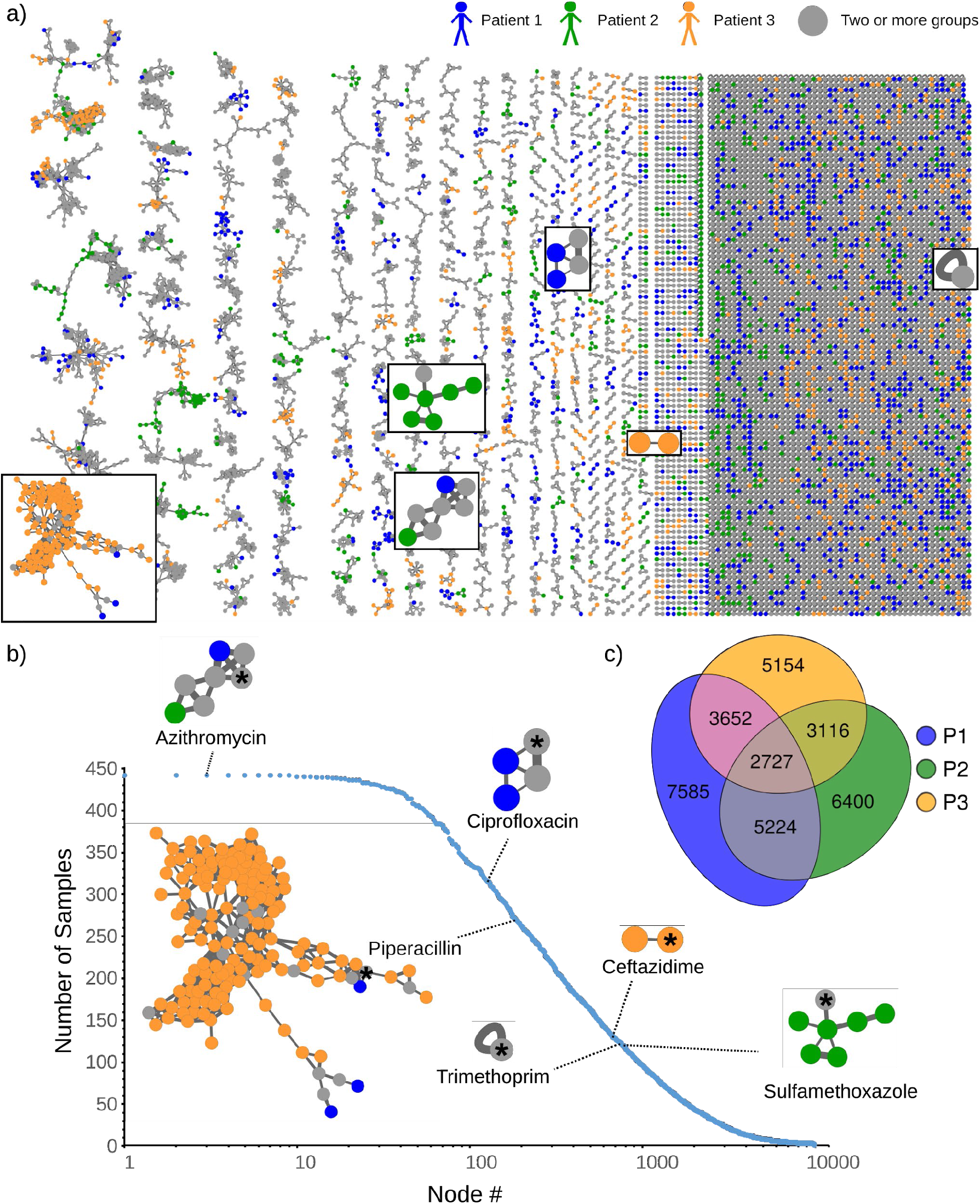
Molecular network analysis of all six lungs from three patients afflicted with CF. a)The molecular network is color coded by patients (patient 1 (blue), patient 2 (green), and patient 3 (orange). The network clusters corresponding to antibiotics are highlighted in boxes. b) The number of samples that contain a given consensus MS/MS spectra (represented as node in Figure 3a) are plotted. The frequency of occurrence of antibiotics detected in this dataset is highlighted on the plot. The number of nodes in a cluster is reflective of detected transformations of the parent compound. The node of parent compound is highlighted by asterisks. The fragmentation patterns of the most frequently observed drug-azithromycin and its analogs are described in Supplementary Figure 7; the large number of nodes in Piperacillin cluster stems from its structural similarity to small peptidic compounds abundant in biological samples and its inherent chemical reactivity with biological molecules (33). c) Venn diagram of the overlap of consensus fragmentation spectra between three patients is shown.

A Procrustes analysis of metabolomics data and 16S rRNA data with closed-reference OTU picking revealed a close association between the microbiome and metabolome in the lung samples (Mantel test *r* statistic = 0.45, *P* < 0.001, *n* = 277) (Supplementary Figure 6 (a,b)). This analysis suggests that the microbial composition of each sample in large part is associated with the corresponding chemical diversity. Additionally, Procrustes analysis performed on metabolomics and 16S rRNA with deblurred sOTUs (34) resulted in the same trend (Supplementary Figure 6 (c,d), Mantel test *r* statistic = 0.38, *P* < 0.001, *n* = 263). A PCoA plot of the metabolome data with Canberra distance showed that a vast chemical diversity exists not only between the patients (Figure 1c), but also within a patient’s own lungs (Supplementary table 2). This suggests that the chemical makeup of the patients with CF disease is highly personalized and that a single CF lung contains unique chemical microenvironments that provide different niches for microbial pathogens to live in. While metabolic diversity between patients in relation to disease state is previously described (32, 35), mechanisms leading to such diversity within the lungs remain poorly understood.

One of the additional benefits of an untargeted metabolomics analysis approach is the ability to track the medications that are taken by the patient, as medical records can oftentimes be incomplete and/or inaccurate due to lack of patient compliance, as well as to identify metabolic transformations of the medications. For example, in the present study, in addition to the prescribed medications from the clinical records (different antibiotics, bronchodilators, two medications for digestive health, medications given during surgery and over-the-counter medications that are used as cough suppressants), antihistamines and multiple over the counter medications have been detected (Table 1). Detailed knowledge of extant exogenous compounds in tissues of interest is important, among other things, to evaluate their effect on the microbiome and microbial interactions for better understanding disease etiology. Another advantage of a molecular networking approach for untargeted metabolomics data analysis is that it allows for postulating structures for unknown compounds, nodes of which are connected to nodes of known compounds (annotation propagation), and is therefore very useful for identifying drug metabolites (Figure 3b). The distributions of drugs and the metabolites can then be evaluated by 3D cartography even in the absence of a stable isotope tracer. Using a molecular networking approach in this study, unknown metabolites that have never been reported before in blood or tissue of humans and animals were detected (Supplementary Figure 7). The unknown metabolite of Azithromycin (*m/z* 382.26) is annotated as methylated-azithromycin where the methylation, based on the analysis of the fragmentation data, occurs in the core macrolide ring structure of azithromycin and another unknown metabolite is proposed to have oxidation in the macrolide ring (Supplementary Figure 7). These modifications of core macrolide structure of azithromycin have not been described previously and their biological activities are unknown. Although these metabolites were not detected in *in vitro* cultures of microbes isolated from these patients in the presence of azithromycin, the possibility that these are microbially-derived warrants further investigation and cannot be ruled out. Specific *in vivo* conditions may be necessary for regulation of microbial genes involved in antimicrobial metabolism.

As with the microbial heterogeneity, we have observed differences in metabolome distributions. Molecular networking and 3D volume cartography of the antibiotics revealed patient-specific metabolism and drug distributions (Figure 4 and Supplementary Figure 8). The distribution of antibiotics was also found to be different between the left and right lungs of the same patient. For example, the antibiotic Piperacillin and its metabolites were abundant in the upper lobes of the right lung of patient 3 but present in relatively lower abundance in the right lung of this patient (Figure 4). In patient 3, there was higher penetration of piperacillin in the upper and middle lobes and poor penetration in the lower lobes of both lungs. Similarly, the antibiotic Linezolid detected in patient 2 had lower relative abundance in the lower lobe of the right lung (Supplementary Figure 8). Overall, the drug metabolites largely follow the same distribution as parent drug except for the glucuronidated metabolite of sulfamethoxazole (Supplementary Figure 8), indicating that metabolism may not be a significant contributing factor for observed uneven distribution of detected antibiotics. Differential vascularization and tissue necrosis also contributes to non-uniform drug penetration in severe end-stage CF disease.

**Figure 4.**
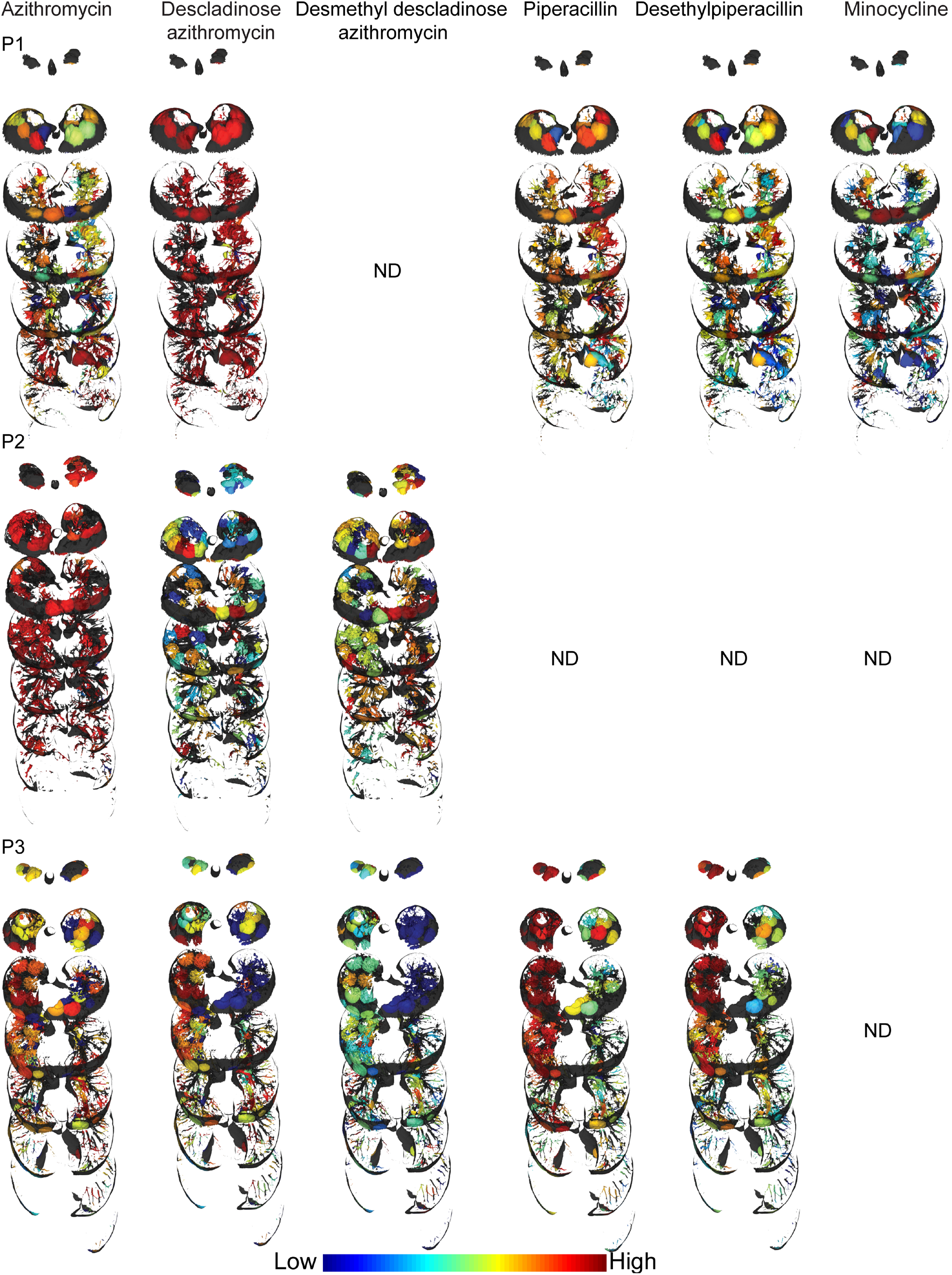
Distribution of selected antibiotics and the metabolites. P1 - patient 1, P2 - patient 2 and P3 - patient 3, ND - not detected. Intensity scale is provided at the bottom (distribution of additional antibiotics and their metabolites is provided in Supplementary Figure 8). Full visualizations of metabolite maps can be accessed via links: <u>patient 1</u>, <u>patient 2</u> and <u>patient 3</u>.

To directly link microbiome and metabolome information and identify associations of compounds detected in the lung tissue with specific microbes, isolates from lungs of all patients were obtained from the clinical lab, cultured directly from swabs of the lung tissue, and MS data was acquired on the organic extracts of the *in vitro* cultures using the same protocol as employed for the tissue extracts. Molecular networking of MS/MS data from culture extracts and tissue extracts provided insights into which molecules are shared between microbes and the human host (Supplementary Figure 9). These molecules include microbe-specific virulence factors, as well as various other molecules such as lipids, fatty acids, amino acid metabolites, dipeptides and tripeptides. Similar to our previously reported observation for one CF lung of a single CF patient (16, 32), a larger diversity of quinolones was detected in cultured isolates as opposed to a smaller diversity of quinolones detected in the lung tissue of all patients dominated by *Pseudomonas* in this study, including a quinolone at *m/z* 268.170 that has never been reported before (Figure 5). Based on MS^1^ and MS^2^ data, the structure of this quinolone is proposed to contain two double bonds in the alkyl side chain as opposed to single double bond found in unsaturated quinolones described in the literature (36) (Supplementary Figure 10). To gain further insight into the variation in the distribution of quinolones in the patients dominated with *Pseudomonas*, we investigated the distribution of quinolones directly within the lungs of these patients (Figure 5b and Supplementary Figure 11). Previously, we reported that the quinolones were prevalent at the upper lobe of the left lung of a single patient (16). In this study quinolones were found to be exclusively present at the upper lobe of lungs of patient 1 and only in the middle of the lungs of patient 3. This indicates that patients dominated by *Pseudomonas* show individualized phenotypes with respect to the expression of these quorum sensing molecules. Furthermore, rhamnolipids, the *Pseudomonas’* biosurfactant, were not detected in the lungs of patient 1 and 3 in this study but were detected in our previous study (16). Patient-specific production of rhamnolipids has been reported previously by culturing isolates in the laboratory but not directly from infected tissue (13). Such compartmentalization of microbial activity within patients, as well as variation between patients is a hallmark of complexity that is inherent to polymicrobial infection in a complex organ; in the present case, a CF lung. Direct visualization of the individual phenotypes in diseased organs enables informed understanding of divergent evolution as well as the spatial molecular environment within a host.

**Figure 5.**
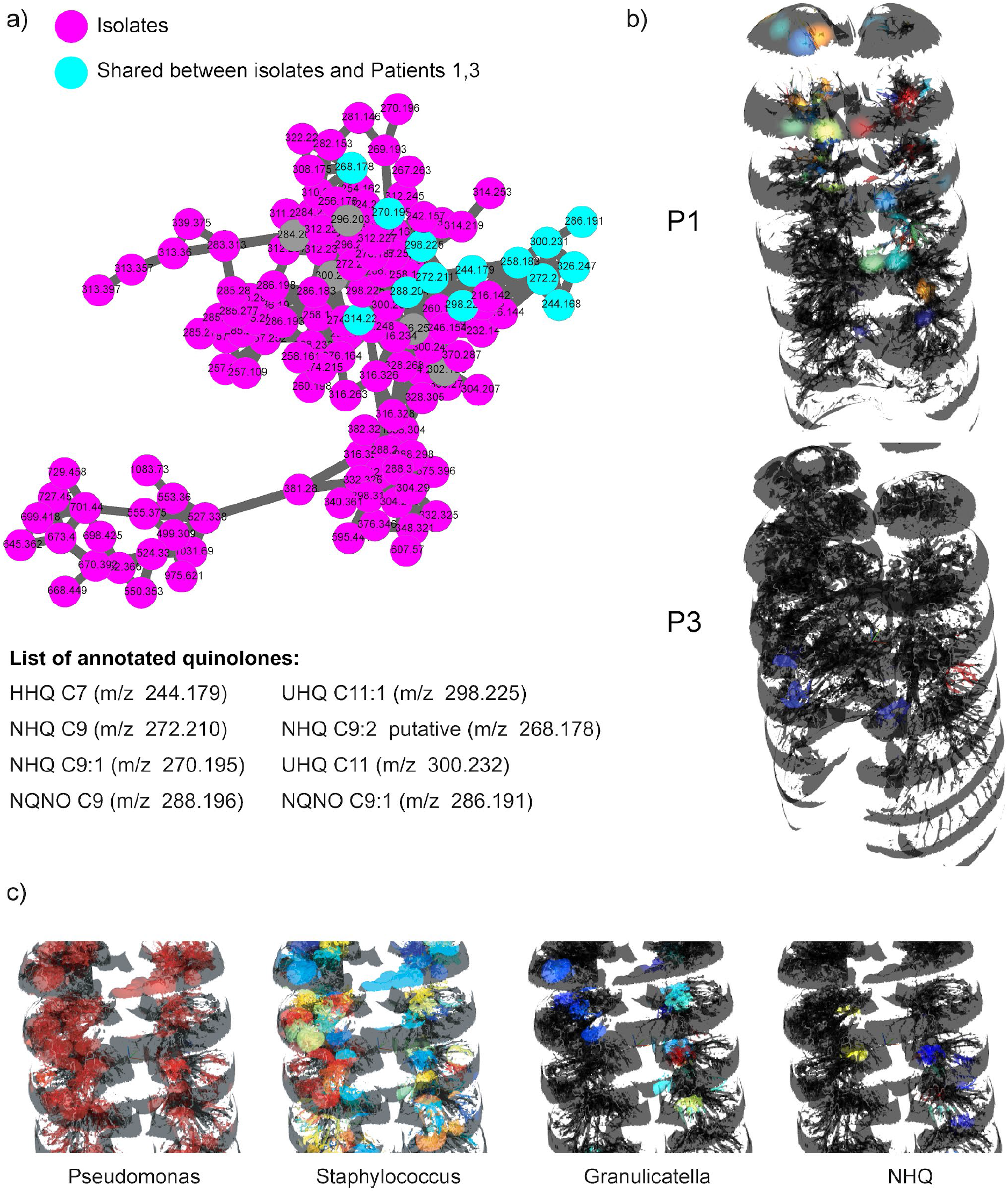
Molecules produced by *P. aeruginosa* in patients 1 and 3. a) The molecular network cluster of quinolones detected in the lung tissue of patient 1 and 3 and *in vitro* microbial cultures of *Pseudomonas* isolated from sputum and the swabs collected from lung sections is shown. b) The distribution of the quinolone HHQ is shown in patients 1 and 3. All the other quinolones showed similar distributions in these patients (Supplementary Figure 11). c) Inset of the distribution of *Pseudomonas*, *Staphylococcus*, *Granulicatella* and the *Pseudomonas* quinolone NHQ in patient 3 suggestive of upregulation in quinolone production by *Pseudomonas* in the regions where interactions of *Pseudomonas* with *Staphylococcus*, *Granulicatella* and possibly other microbes take place. In agreement with this observation, the production of HHQ and NHQ was also found to increase in co-cultures of *Pseudomonas* and *Staphylococcus* compared to Pseudomonas grown alone under identical conditions (Supplementary Figure 12).

The spatial co-distribution of microorganisms, antibiotics and microbial molecules were investigated to establish microbe-metabolite interactions. Although the presence of a single dominant pathogen renders correlation analysis rather uninformative, several trends have been observed. In particular, the distribution of certain microorganisms such as *Staphylococcus* and *Granulicatella* were found to be associated with the distribution of quinolones produced by *Pseudomonas* in patient 3, as shown for NHQ in Figure 5c. We have recently shown that *Staphylococcus aureus* isolated from a CF patient increases quinolone and biofilm production by co-isolated *Pseudomonas in vitro* (37). Similarly, mixing cultures of the *Pseudomonas* and *Staphylococcus* isolated from patient 3 in this study showed increased production of HHQ and NHQ when compared to *Pseudomonas* grown alone under identical conditions (Supplementary Figure 12a). This observation indicates that the production of quinolone molecules is in part also modulated by microbial interactions present in a polymicrobial infection. The complexity of these microbial interactions is further increased as antibiotics cause perturbations of microbial communities reflected by suppression of the virulence factors. Variation in production of quinolones by patient isolates of *Pseudomonas* was observed upon exposure to sub-MIC concentrations (Supplementary Figures 12b). This, and the other observations reported here support the hypothesis that not only are genetic changes responsible for changing metabolism, but microbial interactions in conjunction with multiple other factors including sub-MIC concentrations of antibiotics and perhaps other xenobiotics may also play a role and call for the design of specific studies investigating these phenomena in multiple patient isolates. Thus, it is reasonable to hypothesize that both specific microbial interactions in the lungs and differential abundances of antibiotics could result in metabolic divergence, creating isolated regions of enhanced biofilm formation and tissue damage that is often observed in CF patients by chest X-rays and CT-scans. Application of advanced techniques such as Ultra-High-Resolution Computed Tomography in conjunction with the approach presented here could be a focus of future studies (38).

## Conclusion

Cystic fibrosis is a devastating genetic disease which affects tens of thousands of people worldwide. In this work, we presented the findings of spatial distributions of microbes, medications, and their metabolites throughout lungs of three patients afflicted with CF. We have found that although the microbiome is predominantly patient-specific, the chemical differences between locations within patient’s own lungs may be greater than inter-patient variations. In-depth analyses revealed differential drug penetration, metabolism of prescribed medications, and microbial compartmentalization resulting in metabolic divergence governed by local microbial interactions. Mapping of microbial communities and localized chemistries allowed for visualization of interactions among community members, such as production of quinolones by *Pseudomonas* when present in a community structure with other microbes such as *Staphylococcus* or *Granulicatella.* Visualization of such local infection loci highlight the importance of development of effective drug delivery approaches. Considering recent advances in the development of small-scale robots (as small as few micrometers) that can non-invasively access confined spaces (39), targeted access of internal tissues as well as precision delivery of drug payloads may become feasible in the near future. In general, a paradigm shift of considering localized regions of divergent microbial and chemical distributions is an important next step for effective disease management of polymicrobial infections.

## Supplementary Figures and Tables

**Supplementary Figure 1.**
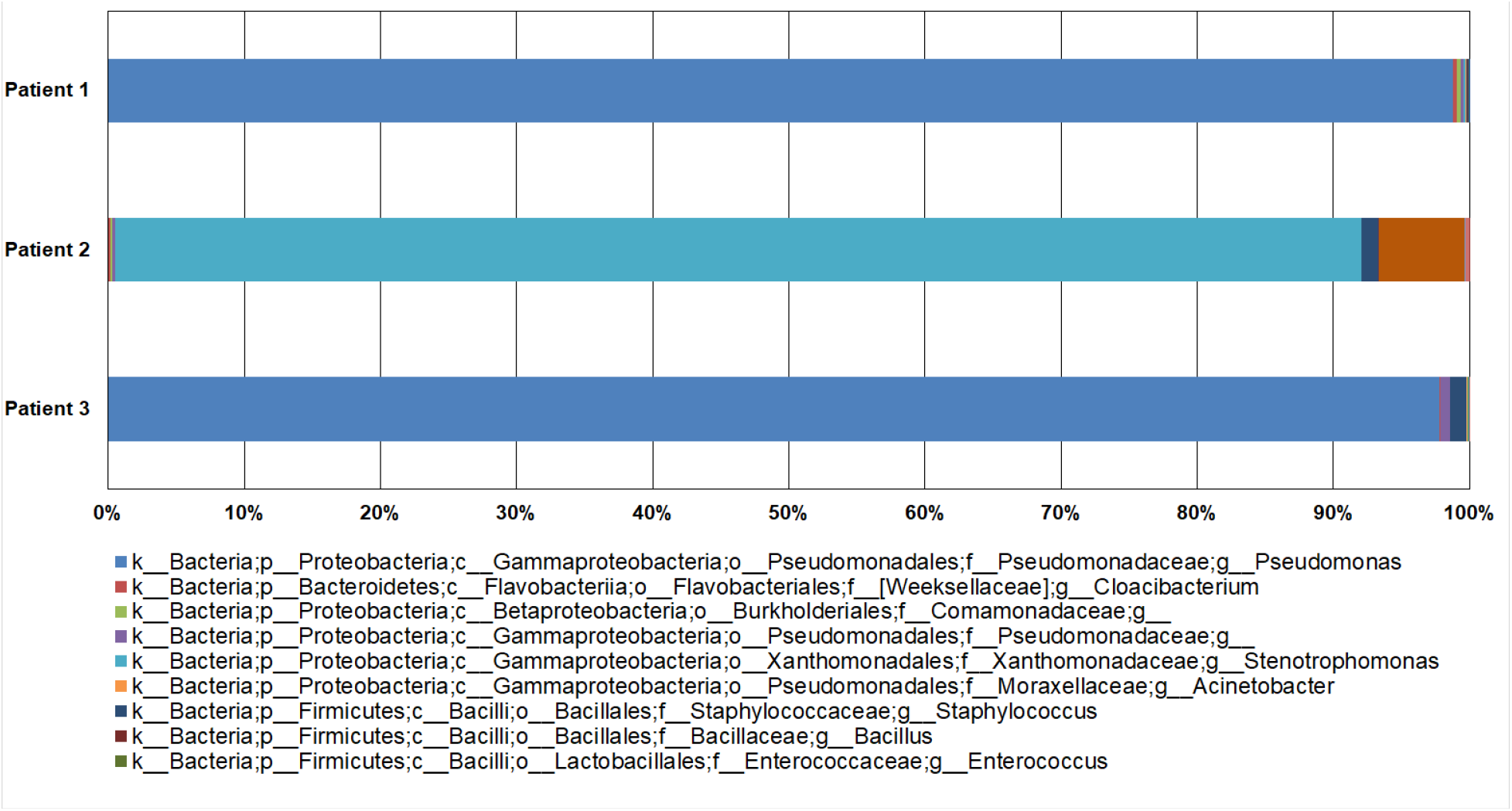
The bar charts of taxonomic summary of all samples for each patient is shown. The legend for 9 most abundant taxa is shown. The complete taxonomic summary is available in Supplementary table 1. The complete description of all OTU’s for each individual sections are available through spatial maps: <u>patient 1</u>, <u>patient 2</u> and <u>patient 3</u>.

**Supplementary Figure 2.**
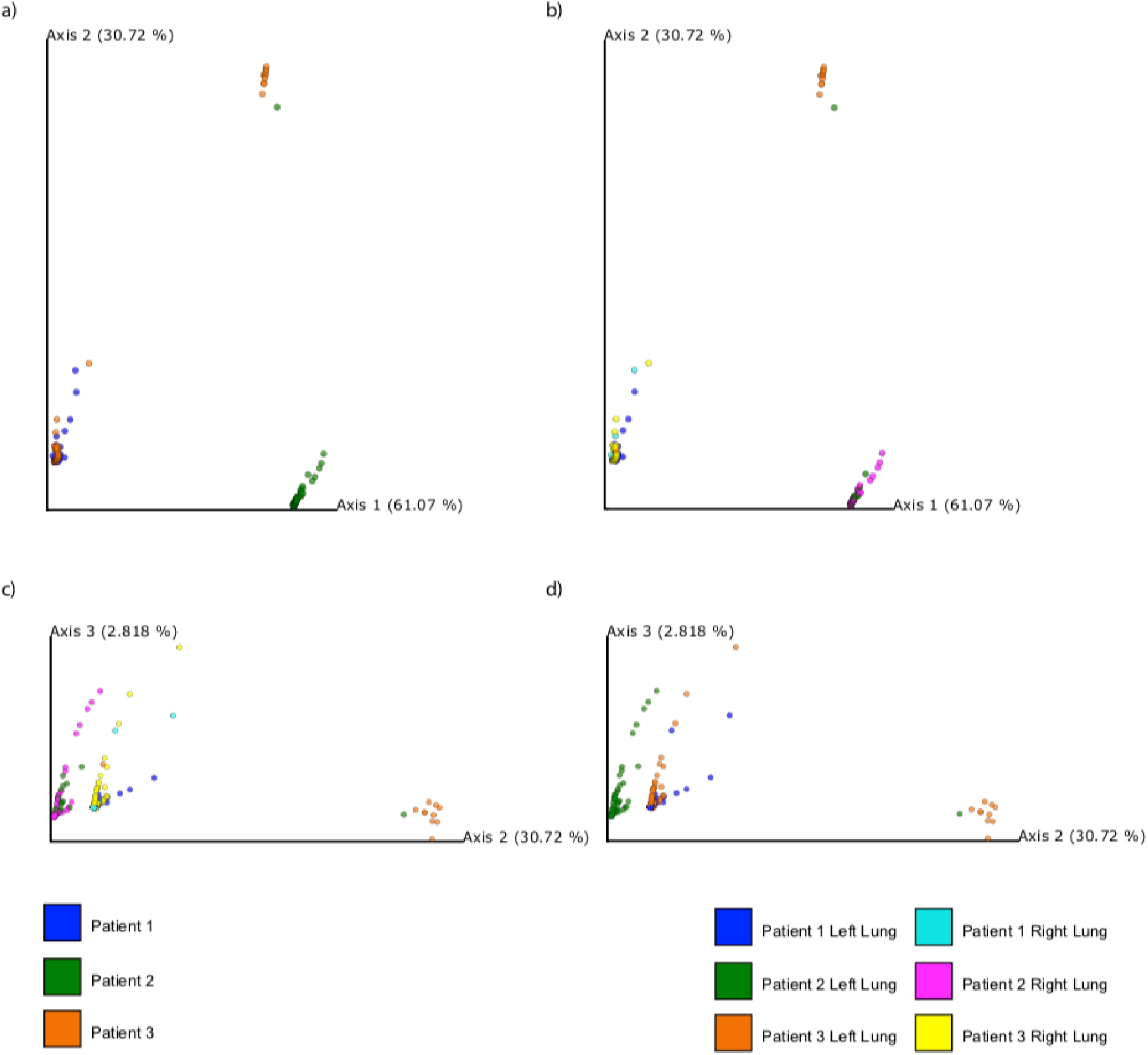
PCoA plot on Weighted Unifrac distance of 16S rRNA data from three patients is shown (40). a) and b) first and second principal component is shown. c) and d) second and third principal component is shown.

**Supplementary Figure 3.**
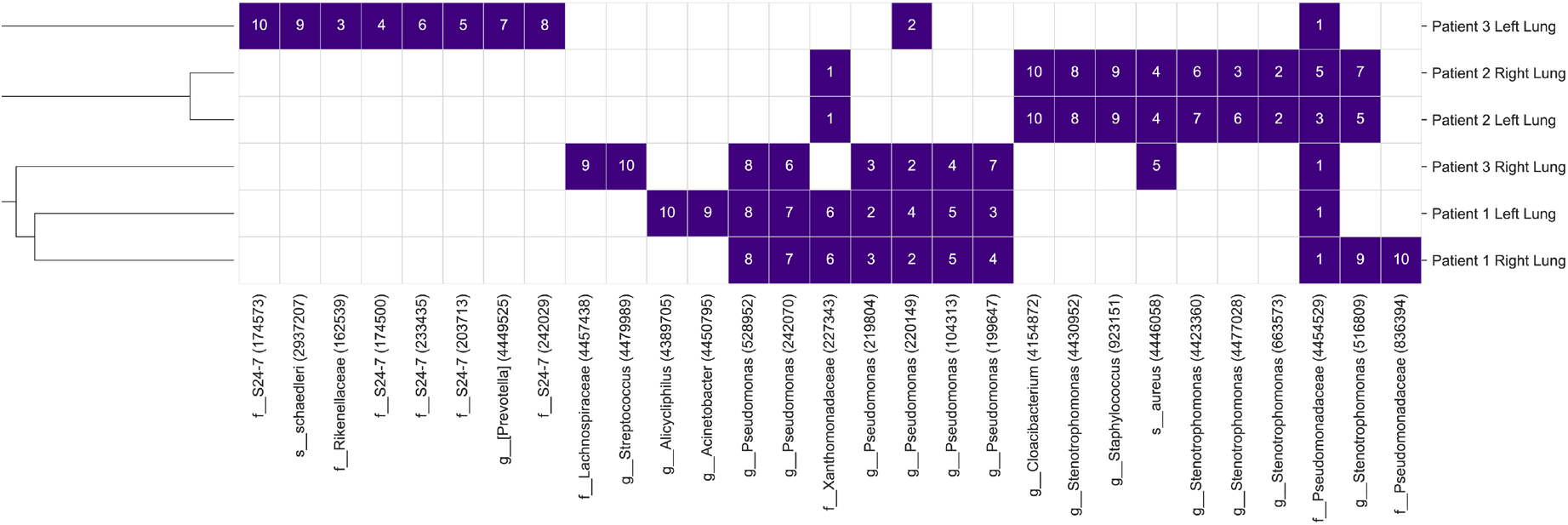
Top 10 most abundant OTUs and their ranks (1 most abundant; 10 least abundant) according to host and sampling site. Columns represent OTUs and are labeled according to their taxonomic classification and their Greengenes identifier. Rows represent the sites and are hierarchically clustered based on the rank vectors.

**Supplementary Figure 4.**
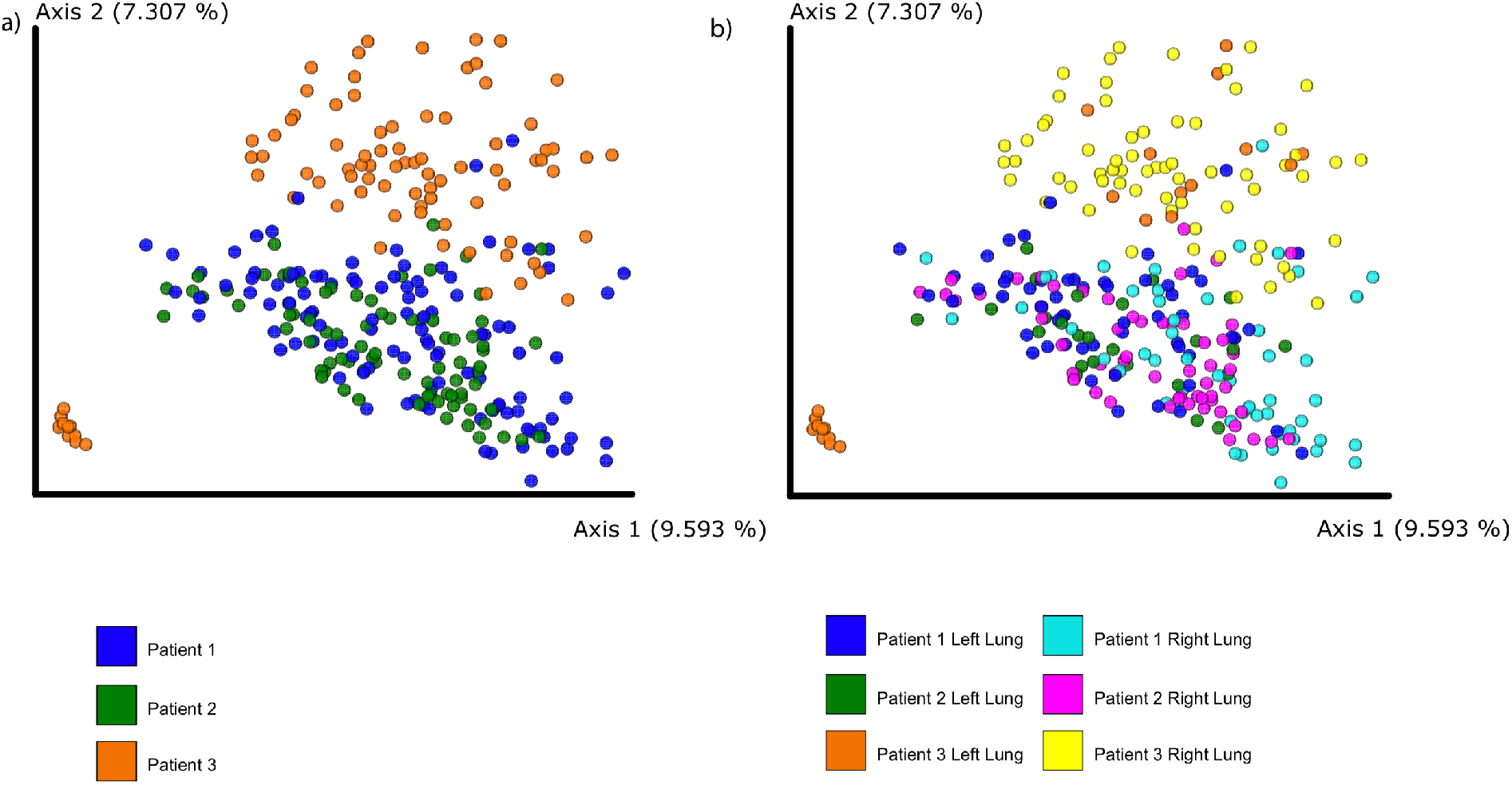
Unweighted UniFrac PCoA plot of the 16S data. a) Ordination colored by subject identifier. b) Ordination colored by subject identifier and sampling site.

**Supplementary Figure 5.**
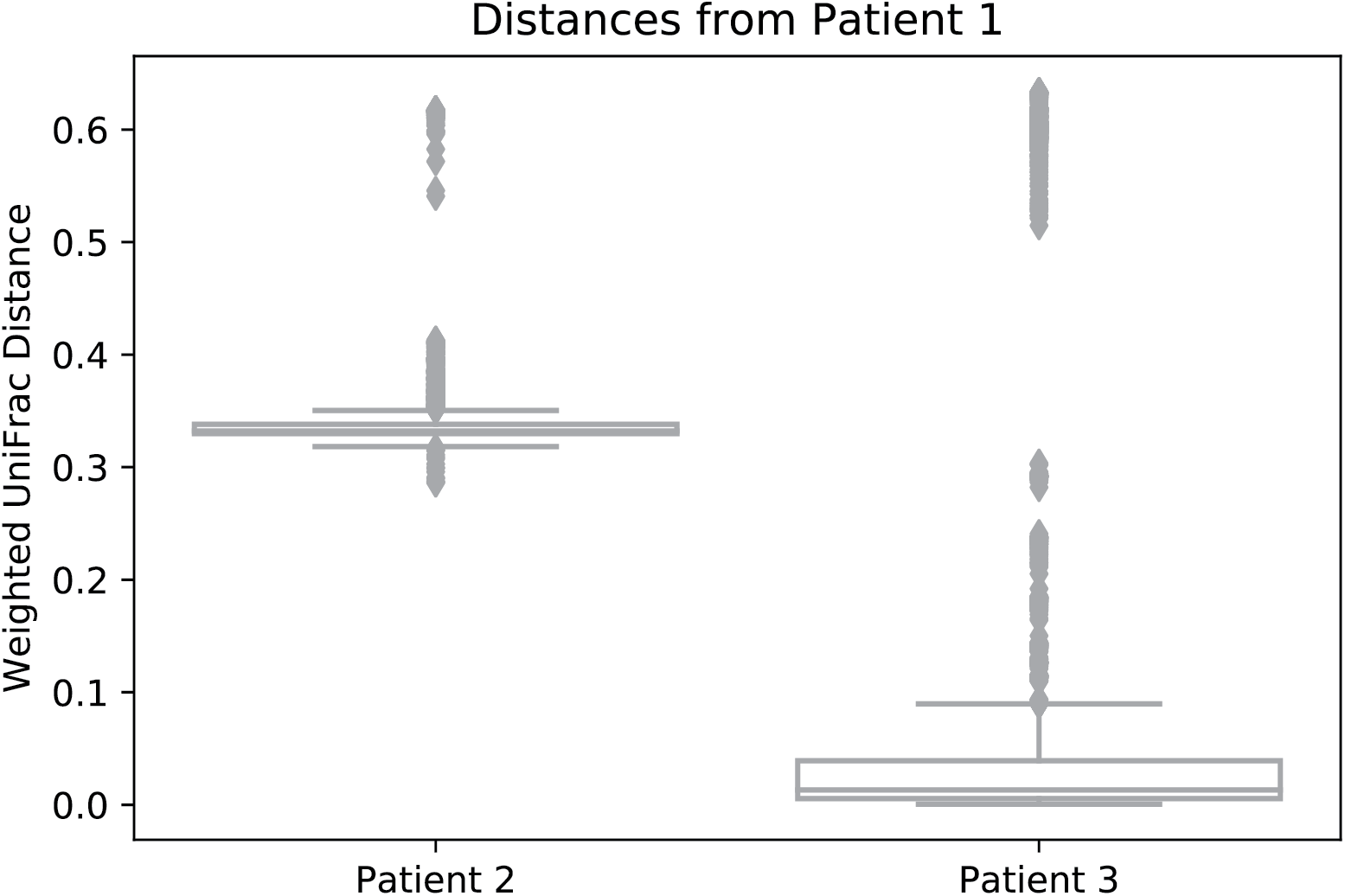
Weighted UniFrac distances from Patient 1 to Patients 2 and 3.

**Supplementary Figure 6.**
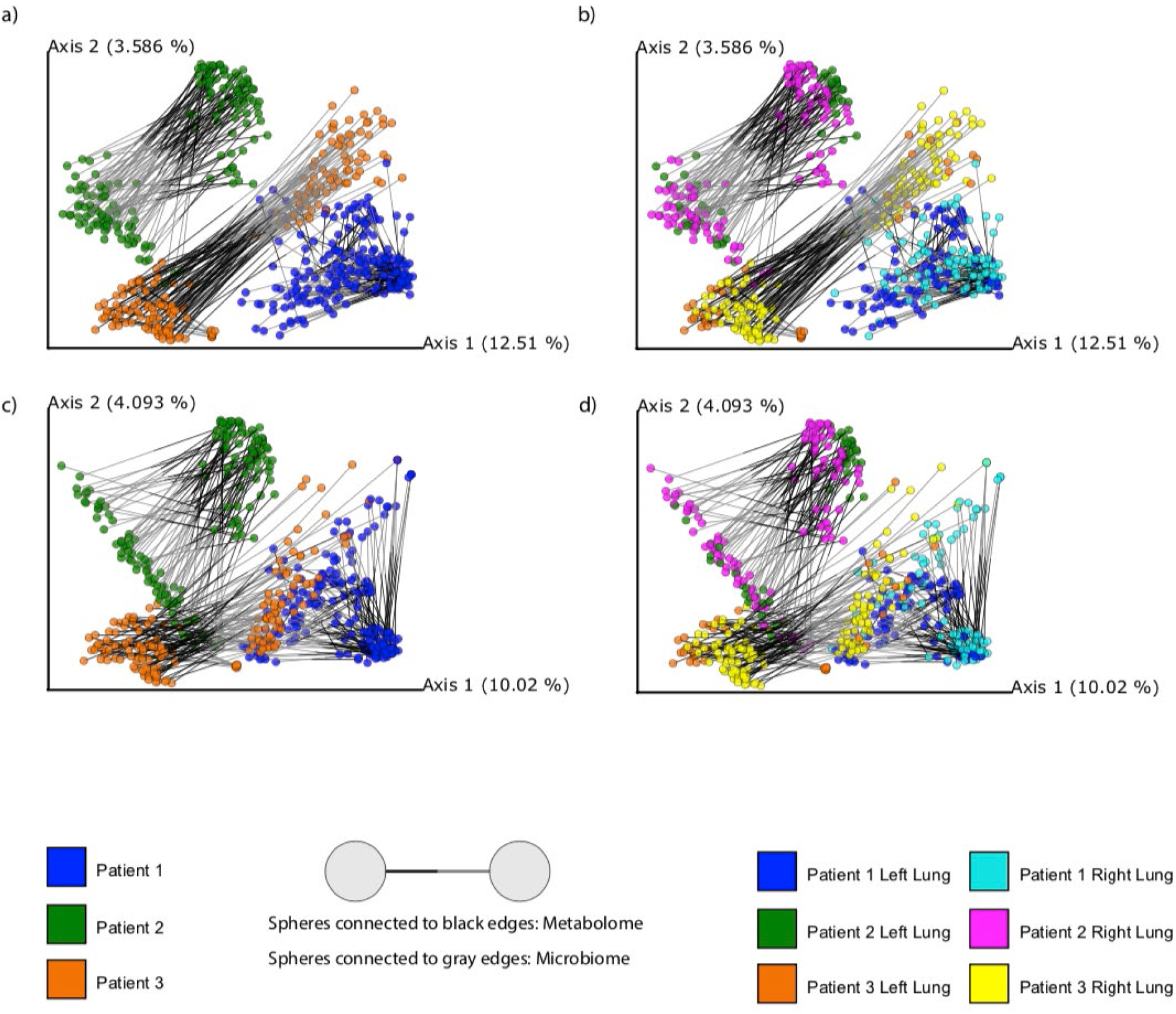
Procrustes plots generated using principal component analysis of Canberra distance metric for untargeted metabolomics data and 16s rRNA sequencing data. a),b) - Emperor visualization plots of metabolomics and closed-reference picked OTUs. c),d) - Emperor visualization plots of metabolomics and deblurred sOTUs.

**Supplementary Figure 7.**
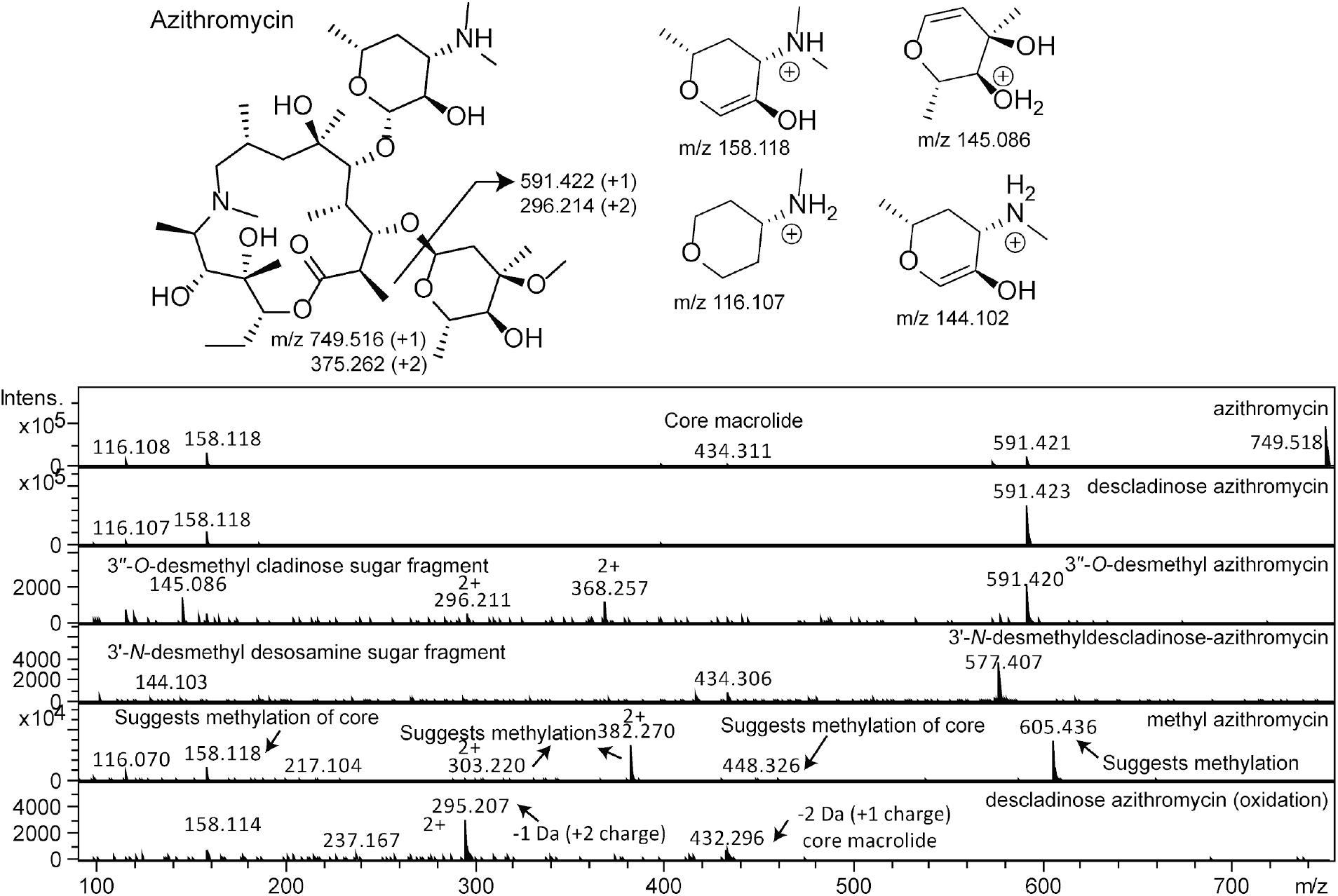
The mass spectral analysis for azithromycin and its proposed metabolites is shown.

**Supplementary Figure 8.**
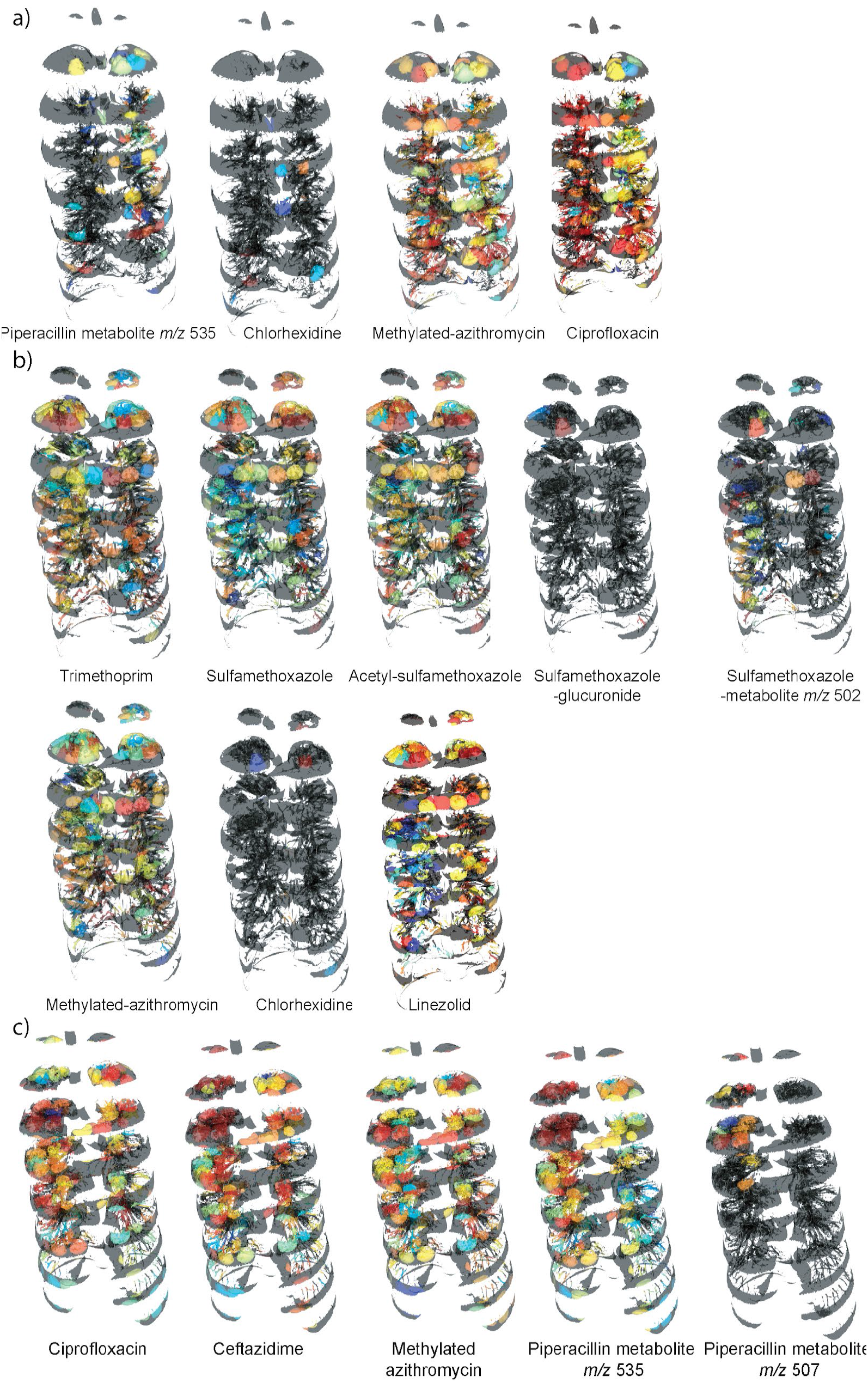
Distribution of antibiotics and its metabolites in patients. a) patient 1, b) patient 2 and c) patient 3.

**Supplementary Figure 9.**
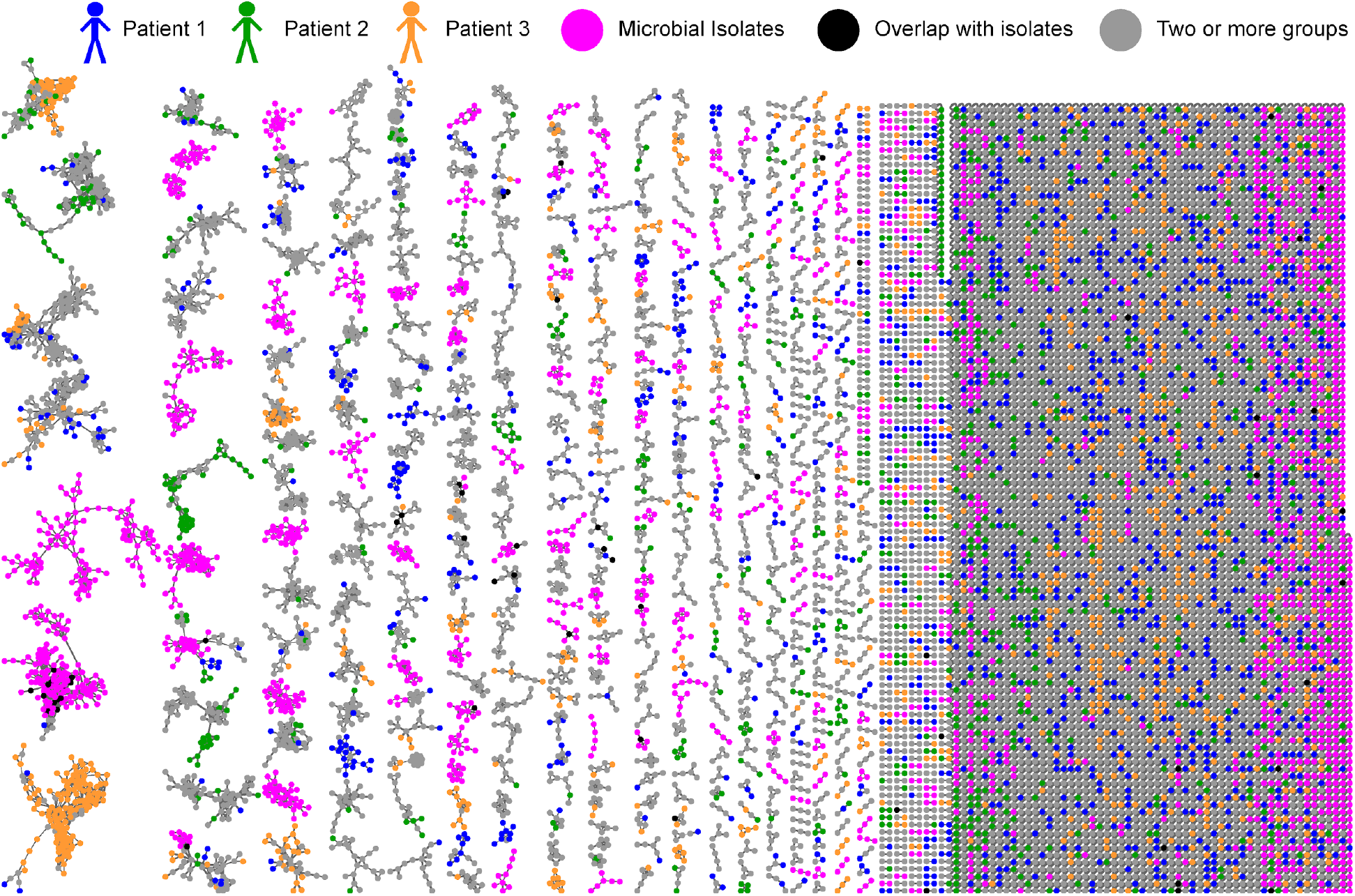
Molecular network analysis of all six lungs from three patients and microbial isolates. The molecular network is color coded by sample source.

**Supplementary Figure 10.**
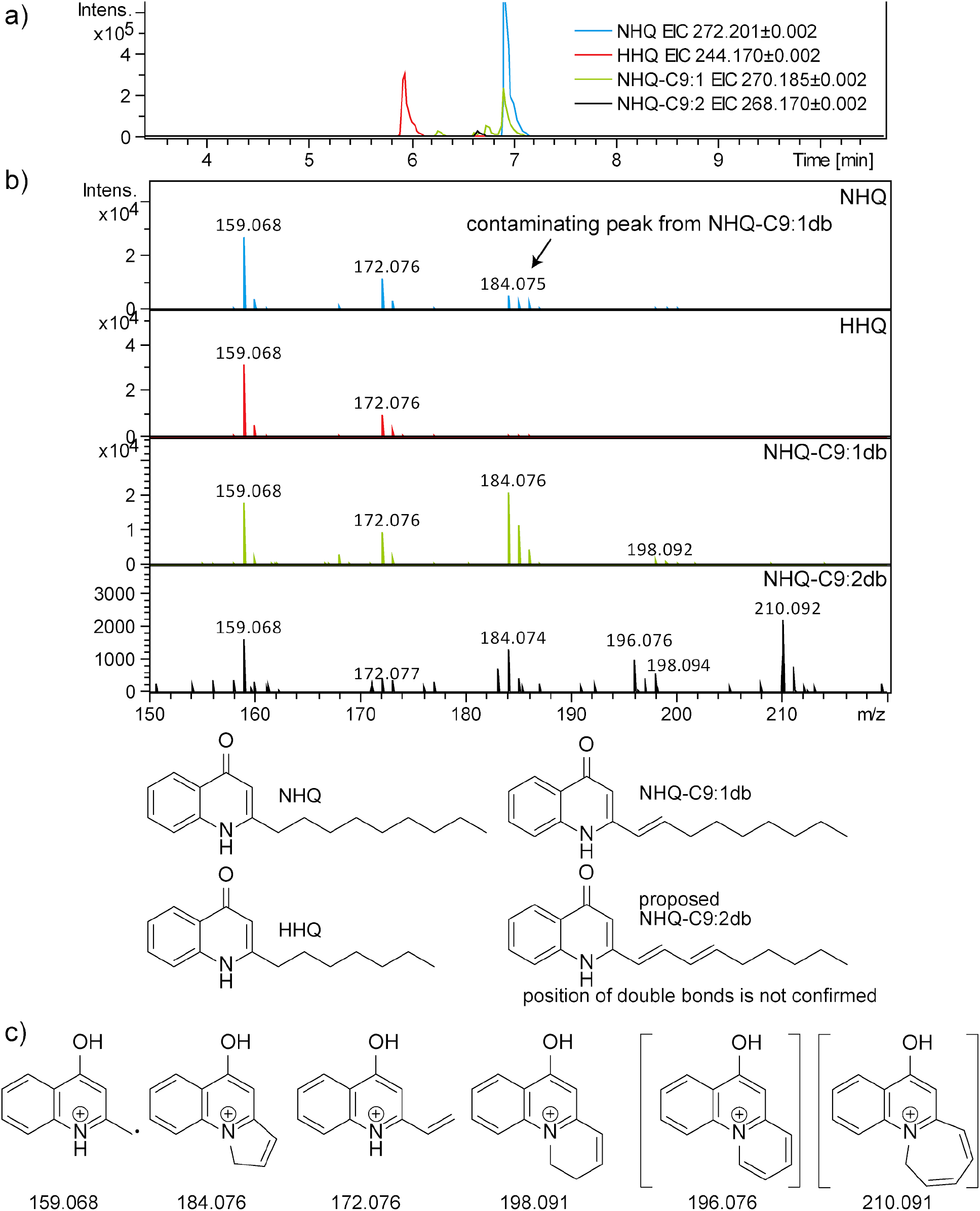
a) The extracted ion chromatograms (EIC), b) mass spectrum and structures of quinolones NHQ, HHQ, NHQ-C9:1db, NHQ-C9:2db (proposed) are shown. The position of additional double bond for NHQ-C9:2db is not known and is putatively drawn. c) The structures of fragments described previously in the literature for NHQ, HHQ and NHQ-C9:1db^40^ and the proposed structures for fragments corresponding to NHQ-C9:2db are enclosed in brackets.

**Supplementary Figure 11.**
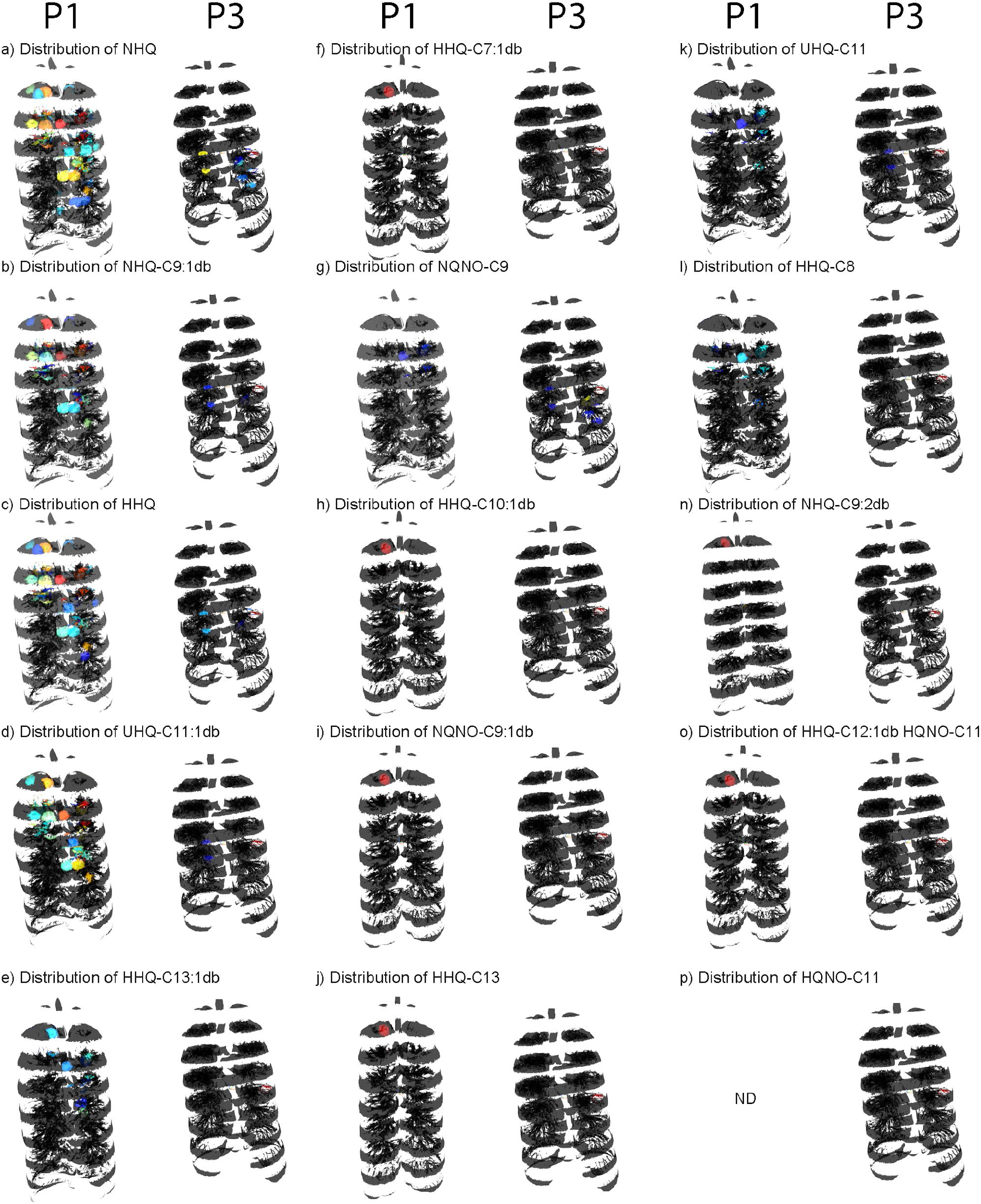
Distribution of quinolones that are common to patients 1 and 3.

**Supplementary Figure 12.**
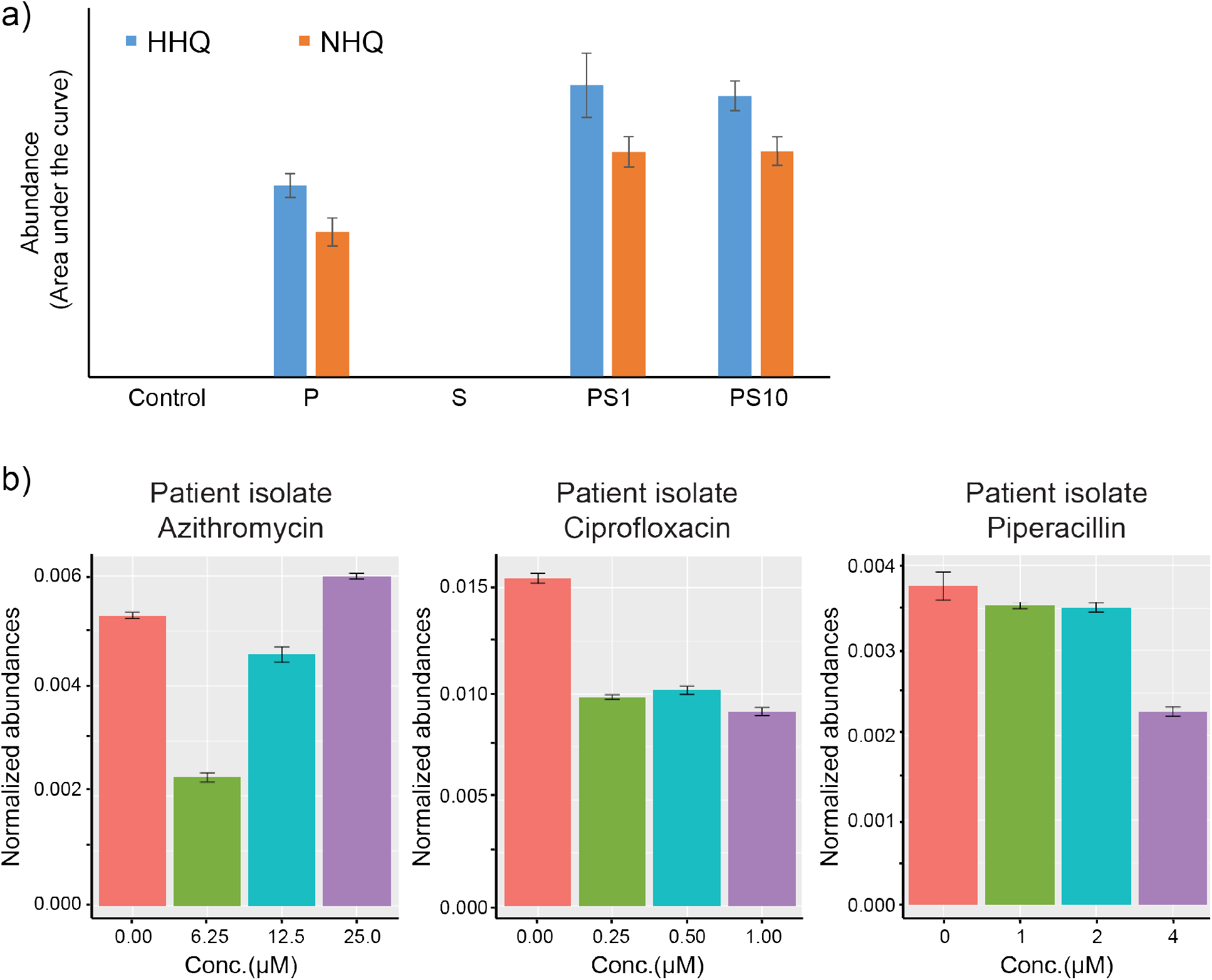
Bar graphs of plots of area under the curves normalized to total ion current (error bars indicate standard deviation from mean values of three independent experiments (*n* = 3)). a) Quinolone production by *Pseudomonas aeruginosa* in single culture and in mixed culture with *Staphylococcus aureus* is shown. PS1 represents cultures were mixed in the ratio 1:1 and PS10 represents a ratio of 1:10, where *S. aureus* culture was diluted 10 fold. b) Production of quinolone NHQ in response to antibiotic exposure by *Pseudomonas aeruginosa* isolated from patient 3. Each color represent different concentrations of antibiotics.

**Supplementary Table 1.**
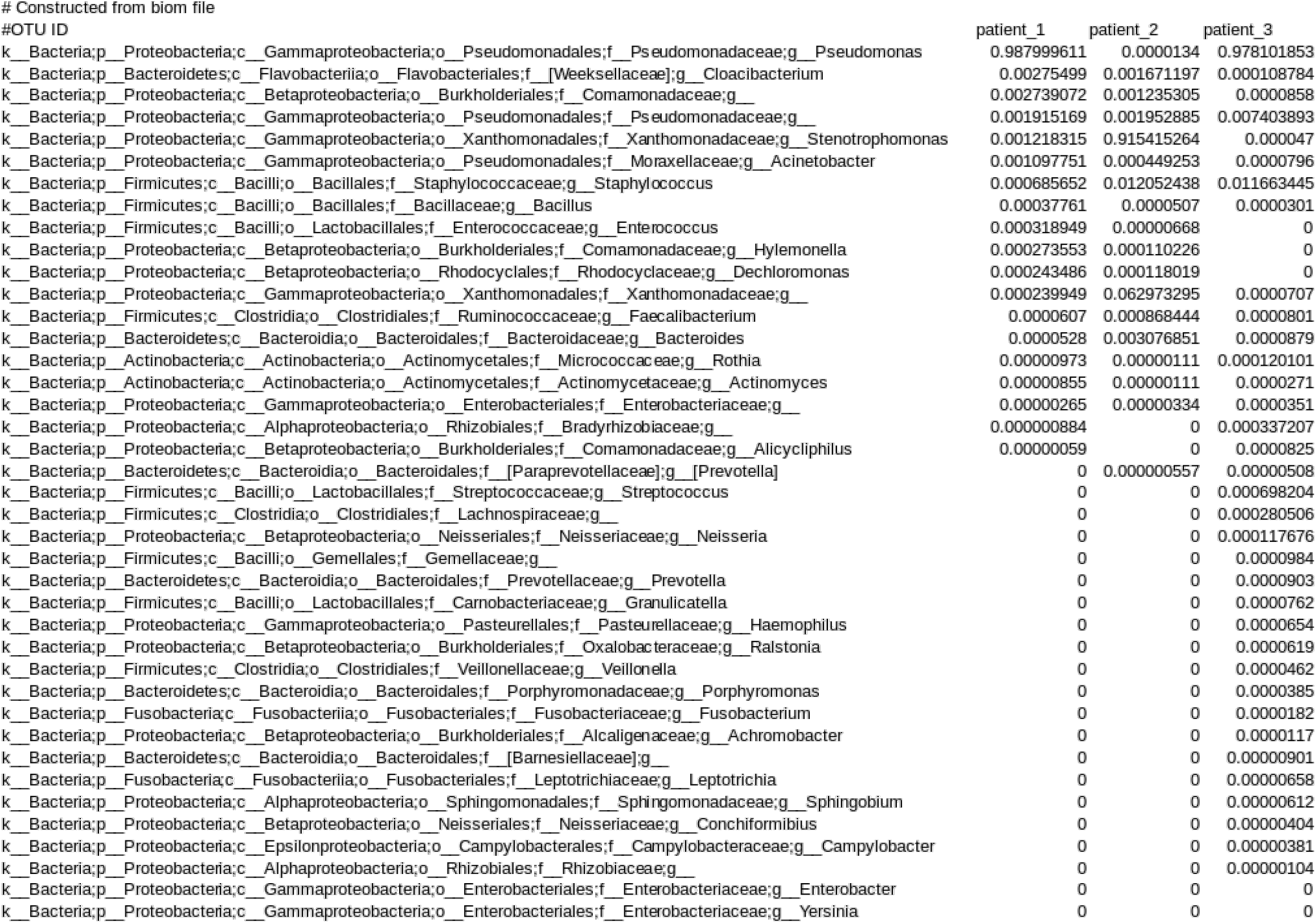
Taxa summary table

**Supplementary Table 2.** Percentage of shared mass spectrometry features between left and right lungs of all patients. This percentage was calculated by taking the features that overlap in comparison to the total number of features associated with each pair of lungs.

|  |  | P1 | P1 | P2 | P2 | P3 | P3 |
| --- | --- | --- | --- | --- | --- | --- | --- |
|  |  | Left | Right | Left | Right | Left | Right |
| P1 | Left | 100 | 88.02 | 75.71 | 76.92 | 70.46 | 72.7 |
| P1 | Right | 88.02 | 100 | 71.67 | 72.39 | 74.01 | 76.73 |
| P2 | Left | 75.71 | 71.67 | 100 | 89.46 | 67.91 | 70.54 |
| P2 | Right | 76.92 | 72.39 | 89.46 | 100 | 70.35 | 72.74 |
| P3 | Left | 70.46 | 74.01 | 67.91 | 70.35 | 100 | 91.56 |
| P3 | Right | 72.7 | 76.73 | 70.54 | 72.74 | 91.56 | 100 |

## Materials and Methods

### Tissue collection and processing

To map the microbiome and metabolome of explanted lungs in 3D, the lungs of three patients were obtained in close coordination with the patient’s physician and the surgical team. This work was approved by the University of California Institutional Review Board (project #081500) and informed consents were obtained prior to tissue collection. The CFTR mutation in Patient 1 was dF508/G551D with no clinical diabetes, in patient 2 was dF508/3120+1G>A with observed clinical diabetes and in patient 3 was dF508/dF508 with observed clinical diabetes. The general workflow for tissue sectioning is described previously (16). Briefly, both the right and left lungs for subject 1, 2, and 3 were collected. The tissue sectioning was performed at the hospital under the guidance of a pathologist. The lungs were first sliced horizontally. The anatomical orientation of each slice was recorded. Every alternate slice starting from apex of the lung was further sub-sectioned into small sections 1-2 cm^3^ in size maintaining the recorded orientation. Each of the sub-sectioned tissue pieces were swabbed with sterile soft foam swabs moistened with Tris-EDTA, pH 7.4. The swabs were stored in 96-well bead plate provided in the PowerSoil®-htp 96 Well Soil DNA Isolation Kit. The plate was placed on dry ice prior and during the collection. The individual tissue pieces were stored in glass jars placed on dry ice. The samples were kept frozen at −80 °C until further processing. Bacterial DNA was isolated from the swabs using the PowerSoil®-htp 96 Well Soil DNA Isolation Kit following the manufacturer’s instructions and was subjected to prokaryotic ribosomal 16S rRNA-based sequencing using the standardized Earth Microbiome Protocol (http://www.earthmicrobiome.org/emp-standard-protocols/). Amplicons were cleaned, pooled and then sequenced on an Illumina MiSeq. Because lungs were obtained at different times, the sequences analyzed for this study were obtained from a total of two individual sequencing runs (sequencing run 1: patients 1-2, sequencing run 2: patient 3). The sequencing runs were performed at the University of California San Diego Institute for Genomic Medicine Center. For untargeted metabolomics analysis, the tissue sections were weighed and extracted with 1 mL/g of tissue with a 2:2:1 mixture of ethyl acetate, methanol and water. An aliquot of 150 µL of the extract was dried for each tissue section and analyzed by MS.

### MS data acquisition

The tissue extracts and extracts of bacterial isolates from the subjects cultured on sheep blood agar and MacConkey agar were resuspended in 80% methanol containing 1µM sulfadimethoxine and analyzed with a UltiMate 3000 UHPLC system (Thermo Scientific) using a Kinetex^TM^ 1.7 µm C18 reversed phase UHPLC column (50 × 2.1 mm) and Maxis Q-TOF mass spectrometer (Bruker Daltonics) equipped with ESI source. The column was equilibrated with 2% solvent B (98% acetonitrile, 0.1% formic acid in LC-MS grade water with solvent A as 0.1% formic acid in water) for 1 min, followed by a linear gradient from 2% B to 100% B in 10 min, held at 100% B for 2.5 min. A small wash segment was employed to wash the column (100% B for 0.5 min, 100%-10 % B in 0.5 min) following which the column was kept at 2% B for A min at a flow rate of 0.5 mL/min throughout the run. MS spectra were acquired in positive ion mode in the range of 50-2000 *m/z*. A mixture of 10 µg/mL of each sulfamethazine, sulfamethizole, sulfachloropyridazine, sulfadimethoxine, amitriptyline, and coumarin-314 was run after every eight injections for quality control. An external calibration with ESI-L Low Concentration Tuning Mix (Agilent technologies) was performed prior to data collection and internal calibrant Hexakis(1H,1H,3H-tertrafluoropropoxy)phosphazene was used throughout the runs. The capillary voltage of 4500 V, nebulizer gas pressure (nitrogen) of 2 bar, ion source temperature of 200 °C, dry gas flow of 9 L/min source temperature, spectral rate of 3 Hz for MS^1^ and 10 Hz for MS^2^ was used. For acquiring MS/MS fragmentation, the 10 most intense ions per MS^1^ were selected and collision induced dissociation energy given in Table 1 was used. Basic stepping function was used to fragment ions at 50% and 125% of the CID calculated for each *m/z* from Table 1 with timing of 50% for each step. Similarly, basic stepping of collision RF of 550 and 800 Vpp with a timing of 50% for each step and transfer time stepping of 57 and 90 µs with a timing of 50% for each step was employed. MS/MS active exclusion parameter was *set* to 3 and released after 30 seconds. The mass of internal calibrant was excluded from the MS/MS list using a mass range of *m/z* 921.5–924.5. The data was deposited in the online repository namely MassIVE and is available under the id MSV000079652 and MSV000079398.

The microbial isolates of *Pseudomonas aeruginosa, Staphylococcus aureus* and *Stenotrophomonas maltophila* from the patients were obtained from the Center of Advanced Clinical Medicine, UC San Diego. The culturing of the isolates and the extractions were performed as described previously (16). The MS data was collected using the same conditions as described above for lung tissue.

### LC-MS/MS data analysis

All mzXML files were cropped with an *m/z* range of 50.00 to 2,000.00 Da and RT range of 0.5 - 18.5 min. Feature extraction was performed using MZmine2 (http://mzmine.sourceforge.net/) with a signal height threshold of 5.0e3 (41). The mass tolerance was set to 10 ppm, and the maximum allowed retention time deviation was set to 0.01 min. For chromatographic deconvolution, the local minimum search algorithm was used with a minimum relative peak height of 1% and a minimum retention time range of 0.01 min. The maximum peak width was set to 1 min. After isotope peak removal, the peak lists of all samples were aligned with the above-mentioned retention time and mass tolerances. After the creation of a feature matrix containing the feature retention times and the exact mass and peak areas of the corresponding extracted ion chromatograms, the metadata of the samples were added. The signal intensities of the features were normalized (probabilistic quotient normalization [PQN]) (42).

Statistical analysis was carried out as follows: QIIME 1.9.1 was used to perform principal-coordinate analysis (PCoA) (beta_diversity.py, a Canberra distance metric in Adkins form). The PCoA plots were visualized in EMPeror (28).

### Molecular networking

The molecular network was created using the online workflow at GNPS platform. The data was then clustered with MS-Cluster with a parent mass tolerance of 0.1 Da and a MS/MS fragment ion tolerance of 0.1 Da to create consensus spectra. Further, consensus spectra that contained less than 3 spectra were discarded. A network was then created where edges were filtered to have a cosine score above 0.7 and more than 4 matched peaks. The edges between two nodes were kept in the network if and only if each of the nodes appeared in each other’s respective top 10 most similar nodes. The spectra in the network were then searched against GNPS’s spectral libraries. All matches kept between network spectra and library spectra were required to have a score above 0.7 and at least 4 matched peaks. The molecular networks and the parameters used are available at the links below:

The molecular network and parameters for the patient data are available at: https://gnps.ucsd.edu/ProteoSAFe/status.jsp?task=6f92a21af31d4569bcdb3cce803c600c

The molecular network and parameters for the patient data and data acquired on cultured microbial isolates are available at: https://gnps.ucsd.edu/ProteoSAFe/status.jsp?task=45d70e56faae4081bbba1f7a9ce38019

In total, 1776 of the nodes were annotated (7.8%) which is higher than the typical rate of annotations of 1.8% annotated in an untargeted metabolomics experiment (30). This is likely because many of the reference MS/MS libraries in the public domain are populated from studies of human samples and also contain most of therapeutics used in the clinic. The error rate of these annotations have been assessed by the GNPS community; with the scoring settings used to obtain the annotations, 1% of which are deemed incorrect, for 4% not enough information is available and 4% could be an isomer or correct, while 91% is presumed correct (30).

### 16S rRNA gene analysis

As described above, sequences were obtained over the course of two months through two independent sequencing runs. The samples for patient 1 and patient 2 were sequenced in one batch and samples for patient 3 were sequenced separately. Each set of sequences were processed and analyzed using Qiita (43). First, the sequencing runs were quality trimmed and filtered using default parameters, resulting in 15,629,914 sequences with a mean length of 150 nucleotides. Next, trimmed sequences (at 150 nucleotides) were clustered into operational taxonomic units (OTUs) using the closed reference OTU picking method at 97% sequence similarity. UCLUST was the underlying clustering algorithm and Greengenes (August 2013 release) was the reference database used (44). This resulted in 340 samples with a mean of 25,078 sequences per sample. After rarefaction at 3,369 sequences per sample 277 samples were used for downstream analyses, including the creation of taxonomy summaries and in the calculation of the UniFrac and Canberra (Adkins form) distances. The most abundant OTU for patient 2 was identified as unclassified genus in the family Xanthomonadaceae. BLAST analysis of the sequence corresponding to this OTU from patient 2 revealed that it belongs to the genus *Stenotrophomonas*. As controls, a total of 49 wells (either containing blank swab or empty well) were interleaved between each of the two sampling sites (left and right lung) of the three subjects. The vast majority of the samples (72%) yielded zero sequences. The remaining 14 samples had a non-zero amount of sequences. Of these, 7 samples were represented by under 14 sequences, a negligible amount compared to the 3,500 sequences per sample used for analysis. And the last 7 samples were represented by over 6,000 sequences each. Although the last sample set was processed without any DNA, the well-to-well contamination during the DNA extraction step yielded these sequences. We removed these samples since the DNA is biological and not representative of a type of actionable contamination (45, 46).

For statistical analysis, QIIME2 (47) was used to perform PCoA (Canberra (in Adkins form), weighted, and unweighted UniFrac distances (weighted UniFrac distances (48)) and Procrustes analysis with metabolomics data. The PCoA and Procrustes plots were visualized in EMPeror (28). The Mantel test was used to calculate r^2^ scores between mass spectrometry and for both closed-reference and deblur 16S rRNA gene analysis data using scikit-bio’s 0.5.5 Mantel’s test implementation.

### 3D lung model generation and visualization

The procedure for creation and visualization of 3D models has been previously described (16). Briefly, the CT-scan images obtained from the radiology department at the Hillcrest hospital in San Diego were combined to create a 3D lung model and exported in the .stl format using InVesalius 3.0. The extraneous pixels corresponding to the chest and back of each model were manually deleted using the 3D modelling software Geomagic Wrap. The relative abundances of detected microbes and molecules were plotted on to these models using a modified version of the ‘ili software available at http://mingwangbeta.ucsd.edu/public/ili/ (16, 49).

### Data availability statement

All data presented in this manuscript is publically available. The metabolomics data was deposited in the online repository namely MassIVE and is available under the id MSV000079652 and MSV000079398. The molecular network analysis and parameters for the patient data are available at: https://gnps.ucsd.edu/ProteoSAFe/status.jsp?task=6f92a21af31d4569bcdb3cce803c600c

The molecular network analysis and parameters for the patient data and data acquired on cultured microbial isolates are available at: https://gnps.ucsd.edu/ProteoSAFe/status.jsp?task=45d70e56faae4081bbba1f7a9ce38019

All raw and processed 16S amplicon sequencing data and metadata are available with Qiita study identification 10169 and as EBI study with accession no. ERP110498.

All figures in this manuscript have associated raw data, which is available through above described accession numbers.

### Code availability Statement

The code for 3D mapping via the browser tool github ‘ili is available at ‘https://github.com/mwang87/ili and the tool is accessed via the link http://mingwangbeta.ucsd.edu/public/ili/

## Acknowledgements

We thank Amnon Amir from the Cancer Research Institute, Sheba Medical Center, Israel for assisting in the analysis of deblurred data used in this article.

